# Anillin-dependent actin assembly at PML NBs protects genome stability

**DOI:** 10.1101/2025.10.26.679380

**Authors:** Vina D. L. Putra, Hannah W. Lay, Caleb Walker, Blake J. E. Lane, Vasiliki J. Willett, Jixuan Gao, Lee L. Marshall, Christopher B. Nelson, Shivam Vora, Milan Sharma, Lucy French, Domonique Jordan, Samuel Rogers, Mark E. Graham, Jacob S. Lewis, Maté Biro, Hilda A. Pickett, Jeanne C. Stachowiak, Anthony J. Cesare, Noa Lamm

## Abstract

Nuclear filamentous actin (F-actin) is emerging as a key regulator of genome stability^1–6^. During replication stress, transient bursts of nuclear actin assembly in S-phase promote fork remodelling and repair^3,4^, but how these bursts are initiated, regulated, and whether nuclear F-actin also act on DNA lesions inherited across cell cycles, remains unknown.

Here, we show that replication stress induces nuclear actin assembly at liquid-liquid phase-separated (LLPS) promyelocytic leukaemia nuclear bodies (PML NBs), driven by the actin-binding protein Anillin. Although best known as a cytokinesis factor, Anillin localises to PML NBs in interphase, where it concentrates monomeric actin (G-actin) and establishes a favourable environment for spontaneous actin assembly. Filament growth within these condensates remodels PML NB morphology and composition, releasing sequestered AKT into the nucleoplasm. AKT is subsequently phosphorylated and activated, and this nuclear AKT activity is required to regulate 53BP1 nuclear bodies in G1. These protective LLPS compartments shield under-replicated DNA inherited from the stressed S-phase until the following S-phase. This prevents premature engagement by aberrant repair pathways that would otherwise generate toxic intermediates and culminate in mitotic failure.

Together, these findings define a condensate-to-filament pathway in which nuclear F-actin dynamically reorganises phase-separated compartments to safeguard genome stability across cell cycles.

## Main

The nucleus is a dynamic compartmentalised environment where biophysical processes such as condensation and liquid-liquid phase separation (LLPS) maintain spatial order in the absence of internal membranes^7–10^. Within the nuclear dynamic space, filamentous actin (F-actin) assembles transiently in response to various stimuli, including serum stimulation^11^, mitotic exit^12^, and genomic insults such as replication stress^3,4^ and DNA double-strand breaks (DSBs)^1,2,6,13,14^. Rapid filament formation and elongation drive dramatic changes in nuclear architecture^3,12,15^, reshaping chromatin organisation^16,17^ and redistributing nuclear bodies^2,3,5,6^. Such structural rearrangements support nuclear functions ranging from transcriptional regulation and splicing^18^ to telomerase recruitment^19^, cell spreading^20^ and the repair of damaged DNA^1,2,6^ and stalled replication forks^3,4^ (reviewed in^21–24^).

During acute DNA replication stress, nuclear F-actin facilitates myosin-dependent repositioning of stalled replication forks to the nuclear periphery, promoting their remodelling^4^ and restart^3^ to maintain genome stability. However, replication stress also impacts subsequent cell cycles through the transmission of under-replicated DNA through mitosis and beyond^25–29^. Whether nuclear F-actin also contributes to managing this prolonged damage response remains unknown.

Moreover, the mechanisms initiating nuclear actin polymerisation during replication stress are poorly defined. It remains unclear whether filaments nucleate stochastically or arise from dedicated nuclear structures. How the biophysical properties of filament growth interact with nuclear condensates to influence signalling and genome maintenance also remains unexplored.

### PML NBs are nucleation hubs for actin

Nuclear F-actin is known to interact with the nuclear periphery to facilitate Homology-Directed Repair (HDR)^2,3,5^. We asked whether nuclear F-actin also engages other nuclear compartments involved in genome maintenance, focusing on promyelocytic leukemia nuclear bodies (PML NBs). PML NBs are LLPS-driven condensates implicated in genome maintenance pathways, including the alternative lengthening of telomeres (ALT) mechanism in telomerase-negative cancer cells^30–33^, and general DNA repair in non-ALT cells^34–37^. They have also been reported to associate with nuclear F-actin^38^.

To examine this potential interaction, we induced replication stress with a mild dose of the DNA polymerase inhibitor Aphidicolin (APH), previously shown to promote nuclear F-actin polymerisation^3–5,19,39^. Super-resolution imaging through the nuclear mid-plane of phalloidin-stained U-2OS osteosarcoma cells, or U-2OS cells expressing a nuclear-localised and GFP-tagged actin chromobody (NLS-actin-CB), revealed short actin filaments emerging from the periphery of PML NBs (Fig. 1a, b).

**Figure 1.**
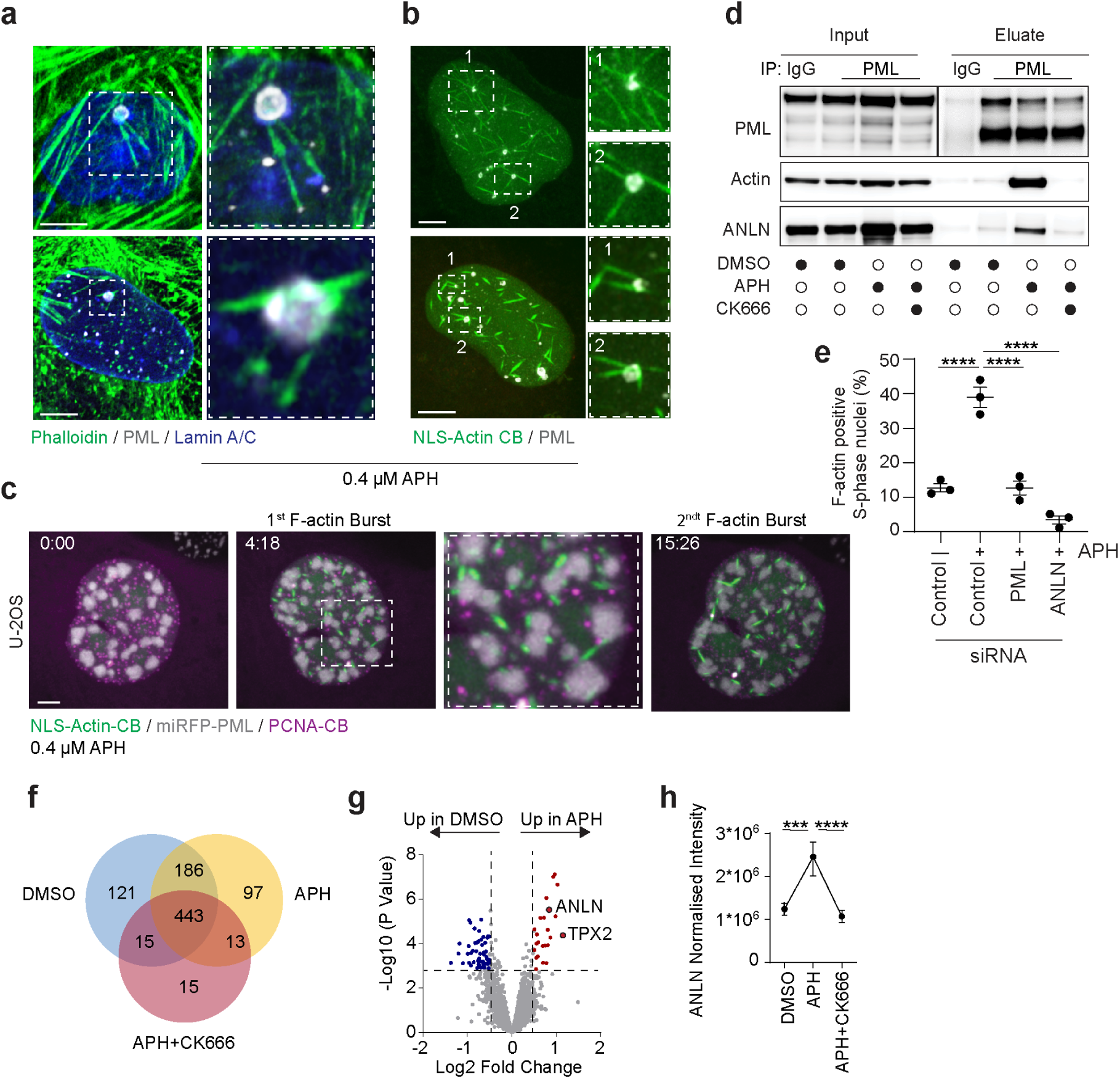
Anillin-dependent nuclear actin assembly at PML NBs. **a**, Representative super-resolution Airyscan microscopy images of U-2OS cells treated with 0.4 µM aphidicolin (APH) for 24 h, showing phalloidin-stained F-actin and PML immunofluorescence. Left: maximum intensity projection; right: Zoomed-in single Z-plane. Scale bar 5µm. **b,** Representative fixed-cell images of U-2OS cells expressing nuclear actin chromobody (NLS-actin-CB) stained with PML antibodies following 24 h of 0.4 µM APH treatment. Scale bar 5µm. **c,** Representative live-cell spinning disk confocal microscopy images of U-2OS cells co-expressing NLS-actin-CB, PCNA-CB, and miRFP670-tagged PML treated with 0.4 µM APH for 24 h. Time is indicated in hours:minutes relative to the first time point. Scale bar 5µm. **d,** Representative immunoblot analysis of endogenous PML immunoprecipitates from U-2OS cells treated for 24 h with DMSO, 0.4 µM APH, or APH in combination with 200 µM CK-666. n = 3 biological replicates. **e,** Quantification of S-phase nuclei positive for nuclear F-actin from experiments in panel (**c**). Cells were transfected with the indicated siRNAs 48 h before imaging and treated with DMSO or APH for 24 h. Data are mean ± s.e.m. n = 3 biological replicates, scoring ≥31 nuclei per replicate. Statistical analysis by one-way ANOVA followed by Tukey’s post hoc test; P < 0.0001 (****). **f,** Venn diagram showing the number and overlap of proteins significantly enriched at PML proximity (significantly enriched at Flag-APEX2-PML vs. Flag-APEX2) in U-2OS cells following treatments described in panel (**d**). **g,** Volcano plots showing enrichment of biotinylated proteins from Flag-APEX2-PML U-2OS cells treated with 24 h of 0.4 µM APH (right) compared to DMSO (left). Significantly enriched PML NB components (log₂ fold change > 0.5, -log p-value > 2.3) are highlighted in red (APH) and blue (DMSO). Statistical analysis by Student’s t-test, n = 3 biological replicates. **h,** Quantification of normalised Anillin (ANLN) intensity from Flag-APEX2-PML U-2OS samples under the conditions described in (**d**). Statistical analysis by one-way ANOVA followed by Tukey’s post hoc test; P < 0.001 (***), P < 0.0001 (****).

To visualise the dynamics between PML NBs and nuclear F-actin, we stably expressed miRFP670-PML and transiently co-expressed nuclear-actin-CB with an RFP-tagged Proliferating Cell Nuclear Antigen chromobody (PCNA-CB). PCNA foci mark S-phase cells. Live confocal imaging over 24 hours revealed two bursts of nuclear actin assembly during S-phase, with filaments consistently emerging from PML NBs (Fig. 1c and Supplementary Movie 1). We observed similar results in telomerase-positive HeLa cells (Supplementary Fig. 1a and Supplementary Movie 2). While visualising nuclear actin filaments is more challenging in HeLa than U-2OS, the data suggest PML NBs serve as nucleation platforms for nuclear actin independent of the telomere maintenance mechanism.

Co-immunoprecipitation (Co-IP) of endogenous PML showed increased actin association following APH-induced replication stress (Fig. 1d). This was suppressed by CK-666, a chemical inhibitor of the ARP2/3 actin branching complex, which is required for nuclear actin assembly following replication stress^2,3,6^ (Fig. 1d) suggesting an interaction between PML, actin and actin regulators.

PML depletion by siRNA reduced nuclear F-actin formation in S-phase cells following APH treatment as quantified by long-term live imaging (Fig. 1e). Notably, PML siRNA did not affect total nuclear actin levels (Supplementary Data Fig. 1b), indicating PML is required for actin assembly rather than nuclear actin import or stability.

### Anillin promotes actin polymerisation at PML NBs

PML NBs harbour diverse factors, including stable core constituents and dynamic, stress-dependent partners^35,37,40^. To identify PML NB-associated factors that potentially impact nuclear F-actin polymerisation, we used unbiased and time-resolved APEX2 interactomics. APEX2 is a promiscuous biotin ligase that covalently labels proteins within ∼10–20 nm in under 60 s^41^.

We stably expressed Flag-APEX2-PML, or Flag-APEX2 as a non-specific interactomics control, in U-2OS and HeLa. Consistent with spatiotemporal precision, we observed punctate streptavidin staining at PML NBs, and enrichment of PML and its interactor DAXX^42^ in streptavidin pulldowns, specifically in Flag-APEX2-PML-expressing cells and only after APEX2 activation (Supplementary Fig. 1c-f).

Flag-APEX2-PML or Flag-APEX2 U-2OS and HeLa cells were treated with DMSO vehicle, APH, or APH and CK-666, then, following APEX2 activation, biotinylated proteins were recovered by streptavidin and identified through LC-MS/MS. As expected, we observed strong PML and DAXX enrichment in all Flag-APEX2-PML samples (Supplementary Fig. 2a, b). As expected, principal component and UpSet analyses showed construct identity (Flag-APEX2-PML vs. Flag-APEX2) was the main driver of variance in recovered proteins across all treatments (Supplementary Fig. 2c, d). We identified 443 and 98 proteins consistently enriched with Flag-APEX2-PML in U-2OS and HeLa, respectively, independent of treatment, representing the core PML NB interactome (Fig. 1f and Supplementary Fig. 2e). The broader network in U-2OS cells likely reflects expanded PML NB functions in this ALT-positive cell line.

Comparing Flag-APEX2-PML-labelled proteins from APH- and DMSO-treated U-2OS and HeLa cells revealed Anillin (ANLN) as the top actin-binding factor enriched under replication stress (Fig. 1g and Supplementary Fig. 2f). Anillin is a conserved scaffold protein essential for cytokinesis and cleavage furrow ingression during mitosis^43,44^, which we prioritised for further study. TPX2 was also enriched, which, like Anillin, is a cytoskeleton-associated mitotic regulator; in this case, a microtubule-binding protein^45^. Both proteins are classically defined as mitotic factors but persist in interphase nuclei, where their non-mitotic functions remain largely unexplored.

Co-IP of endogenous PML confirmed increased Anillin association with replication stress (Fig. 1d), which was lost with CK-666 inhibition of ARP2/3 (Fig. 1d, h and Supplementary Fig. 2g). Functionally, Anillin knockdown markedly reduced nuclear actin filament formation, comparable to PML depletion (Fig. 1e). Notably, total nuclear actin levels remained unchanged with Anillin siRNA (Supplementary Fig. 3a), indicating Anillin specifically promotes nuclear actin assembly rather than regulating nuclear actin content. Anillin thus functions in interphase as a regulator of nuclear actin assembly in response to replication stress.

### Nuclear Anillin is upregulated in response to replication stress

In unperturbed cells, Anillin expression is cell cycle-regulated, rising during S-phase and peaking in mitosis to support cytokinesis^43^. Immunoblotting revealed Anillin is upregulated in the nuclear compartment in response to replication stress in an ATR- and ATM-kinase dependent manner (Supplementary Fig. 3b-d). ATR and ATM, respectively, are master regulators of replication stress^46^, and DSB responses^47^.

To test if increased Anillin levels with replication stress simply reflect APH-induced cell cycle perturbations, we sorted G1- and S-phase cells from a three-colour FUCCI U-2OS reporter line^48^. As expected, baseline Anillin levels were higher in S than in G1. However, when comparing S-phase cells directly, those exposed to replication stress displayed significantly higher Anillin levels than unstressed S-phase cells, and this enhanced upregulation was ARP2/3 dependent (Supplementary Fig. 3e, f). Additionally, PML depletion resulted in a marked reduction of nuclear Anillin levels (Supplementary Fig. 3g). A similar trend was observed for the serine/threonine-protein kinase AKT, which is sequestered at PML NBs to regulate its phosphorylation status^49^, suggesting a role for PML in maintaining its binding partners in the nucleus. Together, the data support a model in which following replication stress, Anillin is upregulated in an ATR-, ATM, and ARP2/3 dependent manner, and retained in the nucleus at PML-NBs.

### Anillin condensates support actin polymerisation *in vitro*

Spontaneous F-actin polymerisation typically requires nucleation factors to overcome kinetic barriers to assembly (reviewed in^50^). PML NB interactomics did not identify actin nucleators such as ARP2/3, formins, or Spire1/2^1–3,6,11,12^, prompting us to test if Anillin could drive actin polymerisation. Phase-separated condensates of actin-binding proteins can facilitate actin polymerisation (reviewed in^51,52^) by presenting a high local concentrating of G-actin and F-actin binding domains promoting spontaneous filament assembly and organisation^53–57^. Anillin contains a central highly disordered region enriched in charged residues, a feature that promotes multivalent electrostatic interactions in LLPS^58^. Supporting this, the N-terminal half of yeast Anillin (Mid1p) has been shown to form phase-separated condensates *in vitro*^59^.

Full-length human Anillin-GFP^60^ purified from insect cells (Supplementary Fig. 4a) formed spherical condensates in conditions commonly used to mimic the crowded intracellular environment (20mM Tris pH 7.4, 150 mM NaCl, 5 mM TCEP, and 3% w/v PEG 8000)^61^. These condensates increased in size and abundance with protein concentration (Supplementary Fig. 4b, c), fused and relaxed into rounded shapes upon contact (Supplementary Fig.4d), and displayed rapid Fluorescence Recovery After Photobleaching (FRAP) (Supplementary Fig. 4e, f), hallmarks consistent with LLPS and dynamic molecular exchange. Consistent with electrostatic interactions driving condensate formation, Anillin condensed without PEG in solutions below 100 mM NaCl (Supplementary Fig. 4g), and partition coefficients for Anillin condensation were higher at lower ionic strengths (Supplementary Fig.4h, i). Finally, Anillin condensate diameters increased as ionic strength decreased (Supplementary Fig. 4j).

To test whether Anillin condensates support actin filament assembly, we examined the ability of purified 5 µM Anillin-GFP to partition 500 nM G-actin monomers in 150 mM NaCl. Latrunculin A (LatA), an inhibitor of actin polymerisation, was included to ensure that partitioning reflected only monomer recruitment and not filament formation. This revealed partitioned G-actin in Anillin condensates with ∼27-fold enrichment (Fig. 2a, b). Adding increasing concentrations of G-actin specifically in experimental buffer lacking LatA, resulted in Anillin condensates transitioning from spherical to elongated, asymmetric structures (Fig. 2c). This coincided with an increased fraction of condensates displaying aspect ratios >1.2 (Fig. 2d) and a higher average aspect ratio across the condensate population (Fig. 2e). These condensate deformations into higher aspect ratio structures, are a recognised signature of filament formation where elongating F-actin exerts protrusive forces on the surrounding matrix^53–57^. Filament formation and condensate deformation were abolished by LatA, confirming that the morphological changes were dependent on actin filament assembly (Fig. 2f). Phalloidin staining selectively labels F-actin. Consistent with actin polymerisation, Phalloidin revealed ring- and rod-like actin bundles within the condensates formed by Anillin and G-actin (Fig. 2f, g), reminiscent of structures observed in VASP and Lamellipodin condensates. Anillin thus forms liquid-like condensates *in vitro* sufficient to drive actin filament assembly. As with other actin-regulating condensates, polymerised filaments within Anillin droplets localise to the periphery, forming rings that deform the condensates into rod-like shapes as polymerisation progresses (Fig. 2g, h).

**Figure 2.**
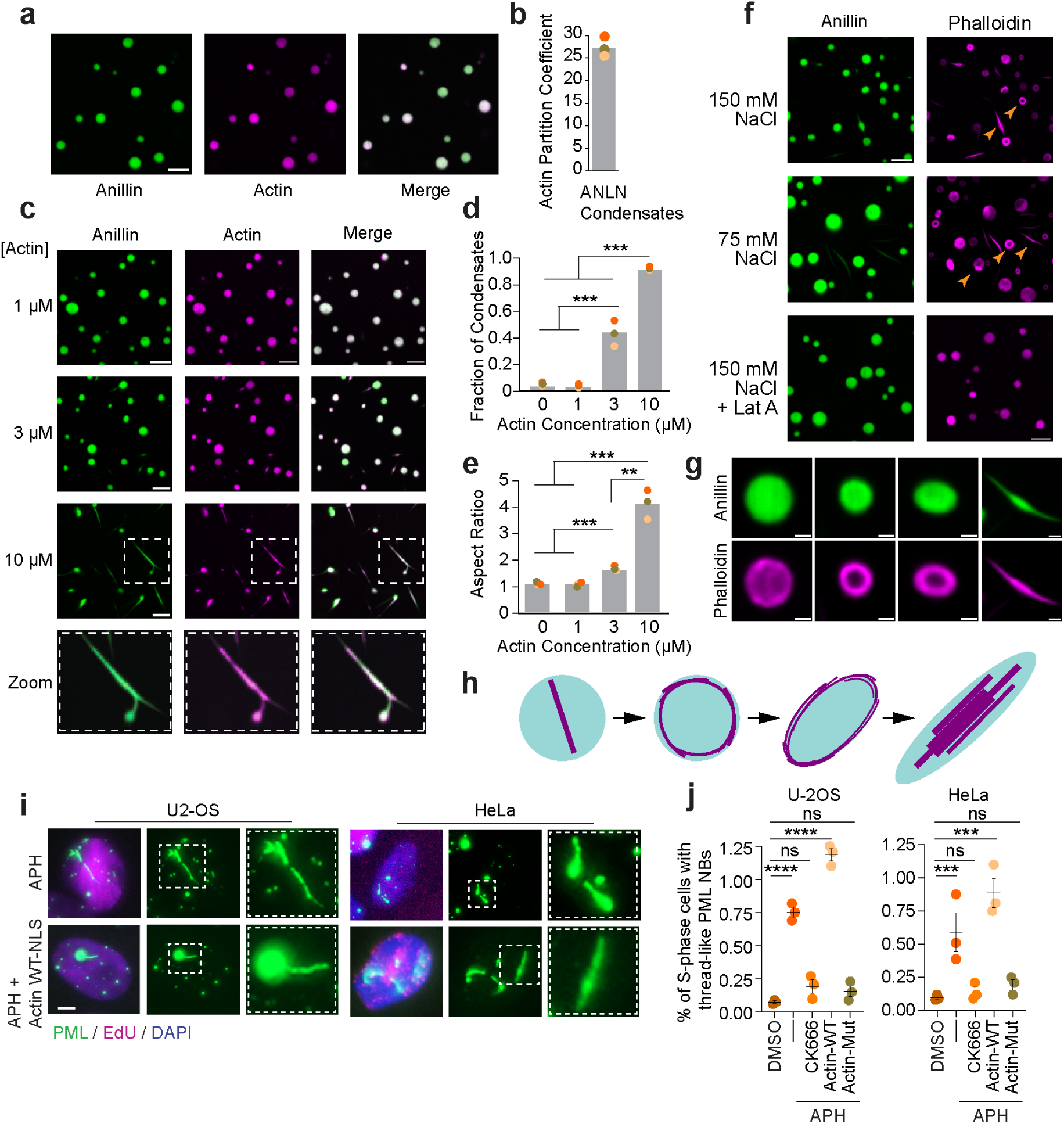
Anillin condensates drive actin assembly and condensate deformation. **a,** Representative spinning disk confocal images showing Atto 594-labelled G-actin partitioning into Anillin condensates (5 μM Anillin). Buffer included 5 μM latrunculin A to prevent F-actin formation. Scale bar, 5 μm. **b,** Quantification of Atto 594-labelled G-actin enrichment within Anillin condensates. The partition coefficient is defined as the ratio of protein intensity inside the condensates to that in the bulk solution. Bar represents the average across three independent experiments. Overlaid circles denote the averages for each replicate. **c,** Representative spinning disk confocal images of G-actin addition at increasing concentrations to Anillin condensates, as in panel (**a**) but without LatA. Bottom panel shows actin filament formation and deformation of condensates at high actin:Anillin ratios. Scale bars, 5 μm. **d,** Quantification of high-aspect-ratio Anillin condensates (aspect ratio >1.2) across conditions shown in (**c**). Data are means of three independent experiments with ≥600 condensates per condition. Overlaid circles represent individual replicate means. Statistical analysis by unpaired, two-tailed t-tests; P < 0.01 (**), P < 0.001 (***). n = 3 biological replicates. **e,** Average aspect ratio of Anillin condensates across the conditions in (**c**), analysed as in (**d**). **f**, Confocal images of Phalloidin-iFluor-594 staining of Anillin condensates incubated with 5 μM unlabeled G-actin. Orange arrowheads indicate actin filaments forming ring and rod structures within condensates under 150 mM NaCl (top) or 75 mM NaCl (middle) buffer conditions. Bottom: addition of 5 μM LatA abolishes actin polymerisation and associated deformation. Scale bar, 5 μm. **g,** Confocal cross-section images illustrating progression of condensate deformation during actin filament assembly. Scale bars, 1 μm. **h,** Schematic model depicting actin assembly within Anillin condensates and the resulting mechanical deformation. **i,** Representative fixed-cell images of U-2OS (left) and HeLa (right) cells with or without transient expression of NLS-Actin-WT, stained for PML and EdU following 24 h of 0.4 µM APH treatment. Insets show thread-like deformation of PML NBs. Scale bars, 5 μm **j,** Quantification of deformed PML condensates in S-phase U-2OS (left) and HeLa (right) cells treated with DMSO, APH (0.4 µM), or APH + CK-666 (200 µM), with or without transient expression of NLS-Actin-WT (Actin-WT) or polymerisation-deficient NLS-Actin-R62D (Actin-Mut). Data are mean ± s.e.m. from three independent experiments with ≥124 nuclei per replicate. Circles show replicate means. Statistical analysis by one-way ANOVA followed by Tukey’s post hoc test; Not significant (ns), P < 0.001 (***), P < 0.0001 (****).

### Actin assembly deforms PML NB’s structure

Actin polymerisation deforms condensates into rod-like shapes as polymerisation progresses (Fig. 2f-h), suggesting similar outcomes may occur within PML NBs upon replication stress. In agreement, rod-like PML NBs were previously reported^62,63^, though the underlying mechanism was unclear. We observed that mild replication stress in U-2OS and HeLa significantly increased ARP2/3-dependent thread-like PML NBs (Fig. 2i, j). Overexpression of polymerisation-competent nuclear actin (3×NLS-Actin^1^ (hereafter nuclear-WT-actin)) further elevated their frequency, whereas a polymerisation-deficient mutant (3×NLS-actinR62D^1^ (hereafter, nuclear-mutant-actin) did not (Fig. 2i, j). We anticipate that the low proportion of deformed PML NBs observed likely reflects the transient nature of filament growth and condensate remodelling, often missed in fixed cell-imaging micrographs. These results indicate that nuclear F-actin polymerisation at PML NBs generates mechanical forces that deform condensates *in cellulo*, paralleling actin assembly-induced deformation of Anillin condensates observed *in vitro*.

### Nuclear F-actin regulates PML NB dynamics

Super-resolution imaging of phalloidin-stained cells revealed frequent associations between extended nuclear actin filaments and PML NBs (Fig. 3a). Live-cell imaging of miRFP670-PML and nuclear-actin-CB expressing U-2OS and HeLa revealed that PML NBs briefly form continuous cylindrical coatings along linear and branched nuclear actin filaments (Fig. 3b and Supplementary Movies 3, 4). Similar “branched” PML NBs were observed in APH-treated fixed cells immunostained for PML (Fig. 3c), indicating this behaviour occurs without exogenous actin or PML manipulation. PML coating of actin filaments was short-lived and reproducibly transitioned into discrete pearl-like droplets decorating the filaments (Fig. 3b, d and Supplementary Movies 3, 4).

**Figure 3.**
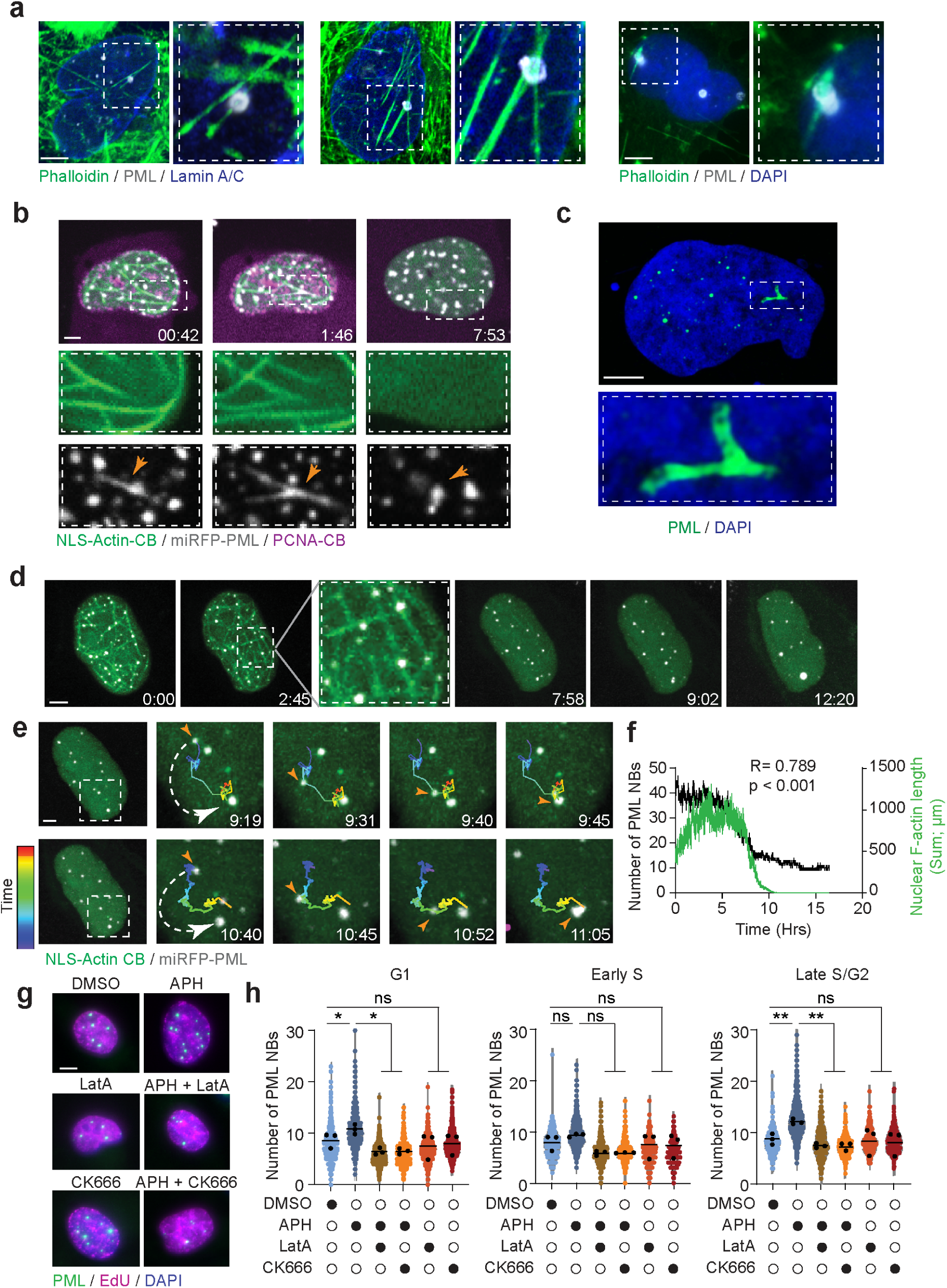
Nuclear F-actin associates with PML NBs and modulates their condensate behaviour. **a,** Representative super-resolution Airyscan microscopy images of U-2OS cells treated with 0.4 µM aphidicolin (APH) for 24 h, stained with phalloidin (F-actin), PML, and either Lamin A/C (left) or DAPI (right). Insets show PML NBs closely associated with nuclear F-actin filaments in a representative Z-plane. Scale bar, 5 μm. **b,** Representative stills from live-cell spinning disk confocal microscopy of U-2OS cells co-expressing NLS-actin-CB, PCNA-CB, and miRFP670-PML, treated with 0.4 µM APH for 24 h. Insets highlight PML NB “wetting” event where PML aligns along branched nuclear actin filaments. Orange arrowhead tracks a representative PML NB over time. Scale bar, 5 μm. **c,** Representative super-resolution Airyscan microscopy image of U-2OS cells treated with 0.4 µM APH for 24 h, stained for PML. The inset shows a branched morphology of endogenous PML NBs. Scale bar, 5 μm. **d,** Representative stills from live-cell spinning disk confocal microscopy of HeLa cells co-expressing NLS-actin-CB and miRFP670-tagged PML, treated with 0.4 µM APH for 24 h. Time is shown as hours:minutes relative to the start of imaging. Inset shows PML NB “pearling” along F-actin. Scale bar, 5 μm. **e,** Time-lapse sequence of two fusion events observed in (**d**). Insets highlight PML NB fusion. White dashed arrows indicate NBs that will fuse; the orange arrowhead tracks the movement of one NB toward another. Trajectories were tracked using Imaris and colour-coded by time. Scale bar, 5 μm. **f,** Line plots showing total length of nuclear actin filaments (green) and total number of PML NBs (black) over time from the movie in (**d**). Pearson correlation analysis was performed across 12 movies from 3 independent experiments, yielding a significant positive correlation (r = 0.789, two-tailed, P < 0.0001). **g,** Fixed-cell images of HeLa cells pulse-labelled with EdU 1 h before fixation and stained for PML following 24 h of treatment with a single drug (DMSO or 0.4 µM APH) or in combination with 200 µM CK-666, or 4 µM latrunculin A (LatA). Scale bar, 5 μm. **i,** Violin plots of PML NB number across cell-cycle phases from (**g**). Black dots represent averages of three independent replicates; solid lines show overall means. Statistical analysis by one-way ANOVA followed by Tukey’s post hoc test; Not significant (ns), P < 0.01 (**), P < 0.001 (***), P < 0.0001 (****).

A likely explanation for this behaviour is wetting, a surface tension-driven process in which liquid-like condensates coat cytoskeletal filaments. Similar interactions occur with Tau-microtubule, TPX2-microtuble^64–68^, and vimentin-actin filaments^69^. Wetting promotes Rayleigh-Plateau instability and fragmentation of a continuous liquid cylinder into discrete droplets (reviewed in^70^), precisely what we observed with PML and nuclear actin fibres. Additionally, time-lapse imaging identified that actin filaments prevented condensate coalescence, with separate PML NBs rapidly fusing once filaments disassembled (Fig. 3d, e and Supplementary Movie 5). We observed a strong positive correlation between total filament length and PML NB number, indicating that F-actin maintains condensates in a dispersed state (Fig. 3f).

We used Quantitative Image-Based Cytometry (QIBC) to measure PML NB abundance in fixed HeLa cultures within a cell cycle context. Mild replication stress conferred G1- and late S/G2-phase increases in PML NBs, abolished by LatA or CK-666 (Fig. 3g, h and Supplementary Fig. 5a). Western blotting confirmed that PML protein levels were unchanged, indicating that the effect reflects altered condensate dynamics rather than protein abundance (Supplementary Fig. 5b). Additionally, APH increased PML NB numbers in HeLa and U-2OS cells expressing nuclear-WT-actin, but not in those expressing the polymerisation-deficient nuclear-mutant-actin (Supplementary Fig. 5c-h). Together, these findings show that nuclear F-actin controls PML NB dynamics through physical alteration of condensate behaviour.

Next, we examined how nuclear F-actin influences PML NB protein composition. Analysis of our interactomics dataset revealed that all PML interactors that were significantly enriched with APH treatment (red dots in Fig. 1g and Supplementary Fig. 2f) exhibited reduced PML interaction with APH plus CK-666 co-treatment (Supplementary Fig. 6a, b, left). Conversely, all interactors exhibiting reduced PML association with APH (blue dots in Fig. 1g and Supplementary Fig. 2f) exhibited increased association with APH and CK-666 co-treatment (Supplementary Fig.6a, b, right). This suggests that both recruitment of new interactors and release of existing components depend on F-actin dynamics.

STRING clustering of APH-enriched proteins revealed that most, including Anillin and TPX2, belonged to mitotic and chromosome segregation pathways (Supplementary Fig. 6c). These same pathways were among the most downregulated with APH and CK-666 co-treatment (Supplementary Fig. 6d), demonstrating that nuclear F-actin actively directs mitotic proteins to PML in response to replication stress.

### F-actin releases AKT from PML NBs

PML NBs sequester nuclear proteins via LLPS^35^. Among these sequestered factors is the serine/threonine kinase AKT, which is retained in an inactive, non-phosphorylated state within PML NBs^49^. The role of AKT in cell survival following genome damage, and as a potential modulator of DNA repair activity^71–75^, piqued our interest. We thus investigated whether nuclear F-actin influences AKT activation, potentially through modulation of PML NB architecture. To do so, we employed a reporter based on the translocation of the AKT substrate FoxO1. Upon phosphorylation by active AKT in the nucleus, FoxO1-mStawberry is exported to the cytoplasm, resulting in a measurable decrease in nuclear fluorescence intensity^76^. Live-cell imaging in FoxO1-mStrawberry U-2OS cells co-expressing nuclear-actin-CB revealed dynamic fluctuations in nuclear FoxO1 levels following APH treatment, with a strongly positive correlation between F-actin polymerisation and nuclear AKT activity (Fig. 4a, b and Supplementary Movie 6).

**Figure 4.**
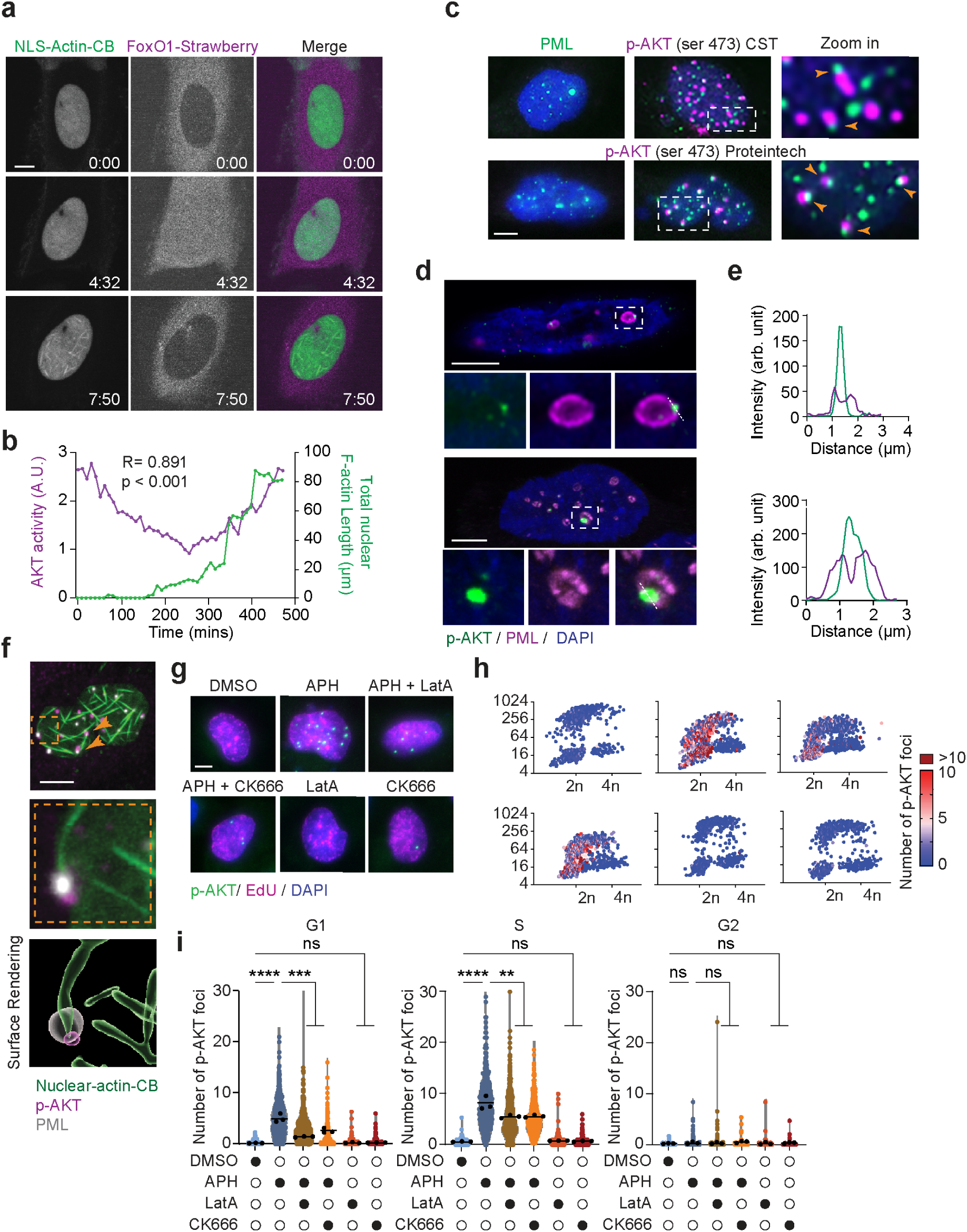
Nuclear F-actin-PML NB association is required for AKT activation. **a**, Still frames from live-cell spinning-disk confocal microscopy of U-2OS cells co-expressing NLS-actin-CB and the AKT activity reporter FoxO1-Strawberry, treated with 0.4 µM APH. Scale bar, 5 µm. **b,** Line plots showing nuclear F-actin filament length (green) and AKT activity (– (FoxO1 nuclear signal); purple) over time from movies in (**a**). Pearson correlation analysis was performed across 15 movies from 3 independent experiments, yielding a significant positive correlation (r = 0.891, two-tailed, P < 0.0001). **c,** Fixed-cell images of U-2OS cells stained for PML and phospho-AKT (Ser473) using CST (top) or Proteintec (bottom) antibodies after 24 h APH treatment. Orange arrowheads mark p-AKT signal adjacent to PML NBs. Scale bar, 5 µm. **d,** Super-resolution Airyscan images of U-2OS cells treated with APH for 24 h, showing single Z-plane staining for p-AKT-Ser473 and PML. Scale bar, 5 µm. **e,** Fluorescence intensity profile along the dashed line in (**d**), showing a p-AKT peak where PML signal decreases. **f,** Airyscan single Z-plane of U-2OS cells expressing NLS-actin-CB after APH treatment, stained for p-AKT and PML. Insets and orange arrowheads highlight p-AKT release from PML NBs at filament ends. Surface rendering by Imaris. Scale bar, 5 µm. **g,** Fixed-cell images of U-2OS cells pulse-labelled with EdU for 1 h before fixation and stained for p-AKT after 24 h treatment with DMSO, APH, or APH combined with CK-666 (200 µM) or LatA (4 µM). Scale bar, 5 µm. **h,** Scatter plot of nuclei from (**g**), showing DAPI versus EdU intensity, colour-coded by number of p-AKT foci. **i,** Violin plots of p-AKT foci across cell-cycle phases from (**g**, **h**). Black dots represent averages of three independent replicates; solid lines show overall means. Statistical analysis by one-way ANOVA followed by Tukey’s post hoc test; Not significant (ns), P < 0.01 (**), P < 0.001 (***), P < 0.0001 (****).

In agreement, immunostaining for pAKT-Ser473 in U-2OS revealed discrete nuclear foci in response to replication stress, the majority of which were in immediate proximity to PML NBs (Fig. 4c). Similar patterns were observed across multiple cell types, including the primary IMR90 fibroblasts, telomerase-positive HeLa, and Saos-2 ALT-positive osteosarcoma cells (Supplementary Fig. 7a, b), indicating that AKT activation at or near PML NBs is conserved in primary and cancerous cells. We note pAKT-Ser473 foci were highest in the two ALT-positive lines, potentially reflecting the expanded functional roles of PML NBs in these cells.

Super-resolution microscopy revealed that pAKT-Ser473 foci at PML NBs emerged at regions of reduced PML signal intensity as indicated by line profile analysis, consistent with local disruptions to condensate morphology (Fig. 4d, e). In agreement, most interactions between pAKT-Ser473 and PML NBs localised at the tips of nuclear actin filaments, suggesting actin-dependent condensate disruption (Fig. 4f and Supplementary Fig. 7c–f). We also observed diffusive pAKT-Ser473 signals emanating from PML NBs, consistent with release of LLPS-sequestered AKT spills into the nucleoplasm before coalescing into discrete foci (Supplementary Fig. 7g).

To determine how inhibiting actin polymerisation impacted nuclear AKT phosphorylation, we used cell cycle-resolved QIBC analysis of pAKT-Ser473 staining. pAKT-Ser473 foci induced after 24 h APH in G1 and early S were reduced with LatA or CK-666 co-treatment (Fig. 4g-i). Notably, PML siRNA significantly reduced nuclear AKT abundance (Supplementary Fig. 3g) and pAKT-Ser473 foci following APH (Supplementary Fig. 7h, i). pAKT-Ser473 foci were unaffected by combined PML knockdown and actin polymerisation inhibition, consistent with an epistatic relationship between F-actin and PML in nuclear AKT activation (Supplementary Fig. 7h, i). Together, the findings suggest nuclear F-actin dynamics promote nuclear AKT activation through its release from PML NBs during replication stress.

### pAKT regulates 53BP1 NBs to protect under-replicated DNA

QIBC analysis demonstrated strong cell cycle dependence of pAKT-Ser473 foci following 24 h APH treatment (Fig. 4h, i). Acute replication stress occurs in S-phase, so it was surprising that pAKT-Ser473 foci peaked in G1 and declined through S. This outcome, however, mirrored 53BP1 nuclear body (53BP1 NB) formation following APH, which also peaks in G1 and early S-phase^25,27,28,77^. 53BP1 is best known as a DNA damage response factor that localises to DSBs to mediate repair^78^. However, it also forms nuclear bodies in G1 at under-replicated DNA inherited from the previous cell cycle. Unlike its canonical role in promoting repair, here 53BP1 protects these regions until they can be resolved in the subsequent S-phase^25,27,28,77^.

Large G1-phase 53BP1 NBs that protect under-replicated DNA co-localise with γH2AX and PML^25^. Co-immunostaining for pAKT-Ser473, 53BP1, γH2AX, and/or PML, revealed strong co-localisation in U-2OS and HeLa, suggesting PML-localised activated AKT is a 53BP1 NB constituent (Fig. 5a, b and Supplementary Fig. 8a). AKT is a collection of three related kinases: AKT1, AKT2 and AKT3. Simultaneous depletion of all three significantly reduced APH-induced 53BP1 foci in G1 and early S, indicative of AKT regulation of 53BP1 NBs (Fig. 5c, d and Supplementary Fig. 8b). We interpret the observed G1 pAKT-Ser473 foci following 24 h of APH as arising in cells that transited mitosis with under-replicated DNA.

**Figure 5.**
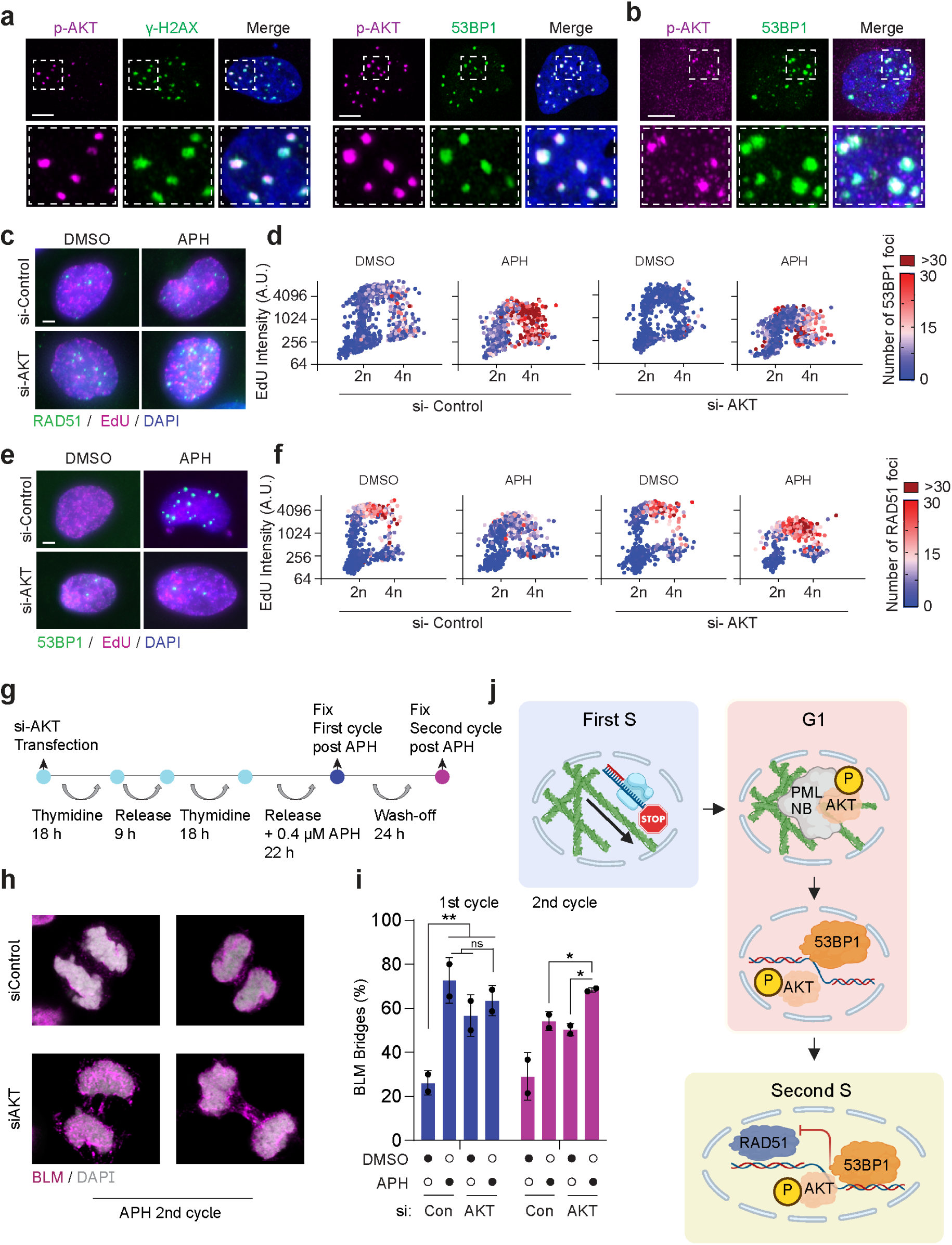
Nuclear AKT is required to protect under-replicated DNA. **a, b,** Airyscan microscopy (single Z-plane) of U-2OS (**a**) or HeLa (**b**) cells treated with 0.4 µM APH for 24 h, stained for p-AKT-Ser473 and either 53BP1 or γH2AX. Scale bar, 5 µm. **c, e,** Fixed-cell images of U-2OS cells transfected with si-Control or si-AKT for 72 h, pulse-labelled with EdU for 1 h, and stained for 53BP1 (**c**) or RAD51 (**e**) following DMSO or 0.4 µM APH treatment. Scale bar, 5 µm. **d, f,** Scatter plots of individual nuclei from (**c, e**), showing DAPI intensity versus EdU incorporation. Each nucleus is colour-coded by 53BP1 (**d**) or RAD51 (**f**) foci number. **g,** Experimental timeline for (**h, i**). HeLa cells were transfected with si-Control or si-AKT, synchronised by double-thymidine block, released into 0.4 µM APH for 22 h and fixed (1st cell cycle) or washed and grown APH-free for 24 h before fixation (2nd cell cycle). **h,** Fixed-cell images of HeLa metaphases stained with BLM and DAPI from the experiment in (**g**). Scale bar, 5 µm. **i,** Quantification of BLM-positive metaphases from (**h**). Graphs show mean values from two biological replicates; black dots represent individual replicates. Statistical analysis by unpaired, two-tailed t-tests; P < 0.05 (*), P < 0.01 (**). **j,** Model of nuclear F-actin in the replication stress response across the cell cycle. In the first replication-stressed S-phase (blue box), nuclear F-actin promotes stalled fork mobility and repair (previously reported^3^). In the subsequent G1 (orange box), nuclear F-actin releases AKT from PML NBs, enabling the formation of 53BP1 nuclear bodies that shield under-replicated regions (this study). In the following S-phase (yellow box), 53BP1 nuclear bodies prevent aberrant RAD51 engagement at under-replicated DNA, thereby safeguarding genome stability (this study).

Inhibiting G1-phase 53BP1 NBs impairs under-replicated DNA protection, leading to aberrant RAD51-mediated repair in the subsequent S-phase^28^. In agreement, combined AKT1, AKT2, and AKT3 knockdown significantly increased RAD51 foci in late S-phase after replication stress (Fig. 5e, f and Supplementary Fig. 8c). Because under-replicated DNA is not a legitimate substrate for RAD51, its inappropriate engagement leads to mis-repair that generates BLM-positive anaphase bridges in the subsequent mitosis^28^.

We thus depleted AKT, synchronised cells, and completed one APH-treated cycle before fixation or allowance of a second APH-free cell cycle (Fig. 5g). Anaphase bridges after the first cell cycle reflect direct replication stress. In contrast, those in the second APH-free cell cycle represent mismanagement of under-replicated DNA. AKT depletion significantly increased the fraction of BLM-positive bridges in the second cycle (Fig. 5h, i), signifying a role for AKT in protecting inherited under-replicated DNA in G1.

We also investigated how AKT influences replication stress survival. Inhibiting AKT with the selective inhibitor MK-2206^79^ markedly enhanced APH-induced growth suppression (Supplementary Fig. 8d), and induced cell death with replication stress as measured by colony formation assay (Supplementary Fig. 8e, f). Additionally, MK-2206 significantly reduced the IC₅₀ of the replication stress-inducing chemotherapy Hydroxyurea (HU), consistent with activated AKT conferring replication stress resilience (Supplementary Fig. 8g, h).

### Anillin is required to establish 53BP1 NBs

Together, our findings support a model where nuclear F-actin releases AKT from PML NBs, resulting in its activation. This AKT activity is required for G1 phase 53BP1 NB formation at under-replicated DNA inherited from the prior cell cycle. As we (Supplementary Fig. 8a) and others have observed, under-replicated DNA in G1 is protected within LLPS structures containing both PML and 53BP1^25^. How nuclear F-actin relates to these G1 NBs was unclear.

Our prior analysis using live imaging to measure nuclear F-actin polymerisation focused on the S-phase during acute replication stress and did not examine the following G1^3^. Consistent with prior work^3,4,39^, we observed here that acute replication stress elevated nuclear F-actin frequency during S-phase (Fig. 1e and Supplementary Fig. 9a). Strikingly, we observed nuclear F-actin in G1, but only when it followed a prior replication stressed cell cycle (Supplementary Fig. 9a). Additionally, these G1-phase nuclear actin filaments that followed a completed cell cycle with replication stress were Anillin dependent (Supplementary Fig. 9a). This is consistent with our above observation of elevated Anillin protein levels in the subsequent G1 after cell cycle completion with replication stress (Supplementary Fig. 3f).

Additionally, we analysed G1 pAKT-Ser473 relative to Anillin function. Consistent with its known role in cell division, Anillin siRNA caused cytokinesis failure^43^, complicating cell cycle resolved analysis. However, total foci counts revealed a significant reduction in pAKT-Ser473 foci with Anillin depletion consistent with the PML-Anillin axis regulating nuclear F-action polymerisation as described above (Supplementary Fig. 9b, c). Moreover, Anillin depletion suppressed 53BP1 NBs (Supplementary Fig. 9d, e). Together, these findings demonstrate that Anillin is required for nuclear F-actin assembly in G1 following replication stress, enabling p-AKT-Ser473 foci formation and establishing 53BP1 NBs (Model; Fig. 5J).

## Discussion

This study revealed that nuclear actin assembled specifically at PML NBs in response to replication stress (Model; Supplementary Fig. 9f). We identified Anillin, an actin-binding mitotic regulator that persists in the nucleus during interphase, as a key driver of this process. While Anillin’s mitotic functions are well established, its nuclear role has remained unclear. We showed that replication stress induced ATR- and ATM-dependent upregulation of nuclear Anillin, which became enriched at PML NBs to promote actin assembly. Notably, we uncovered tight regulation of Anillin by the ARP2/3 complex, both at the level of protein expression and PML association (Fig. 1d and Supplementary Fig. 3f). However, whether this reflects indirect control through branched actin dynamics or a direct effect on Anillin is still unknown.

Although Anillin is not a classical nucleator, our *in vitro* droplet assays demonstrated that Anillin condensates partitioned G-actin and created an environment that supported spontaneous filament assembly. Polymerisation within these condensates generated mechanical forces that deformed them, providing a mechanistic explanation for the enigmatic thread-like PML NB morphologies reported in earlier studies^62,63^. Importantly, we detected these structures in unmanipulated, fixed cells, providing *in vivo* evidence that nuclear actin actively modulated LLPS-driven condensates. This model also explains the pulsatile nature of nuclear F-actin: filaments emerge from PML NB condensates, disrupt the phase-separated state that supports further polymerisation, and terminate growth until condensate reformation enables the next burst.

The interaction between nuclear F-actin and PML NBs extended beyond nucleation. Live-cell and super-resolution imaging revealed dynamic contacts consistent with wetting, a surface tension-driven phenomenon where condensates coated cytoskeletal filaments. This process produced alternating phases in which PML NBs spread as cylindrical sheaths along filaments or reorganised into multiple pearled foci, resembling dewdrops on a spider web. Such wetting-driven remodelling constrained condensate size and prevented uncontrolled coalescence. Complementary proteomics showed that these F-actin-dependent structural dynamics reprogrammed PML NB’s interactome under replication stress, underscoring their functional significance.

Crucially, we demonstrated that nuclear F-actin-PML NB interactions released sequestered AKT into the nucleoplasm, where it became phosphorylated and activated (Model; Fig. 5j). This revealed a unique mode of protein activation through condensate-filament interplay. Activated AKT was subsequently incorporated into 53BP1 NBs. These nuclear bodies protect under-replicated DNA inherited from the stressed S-phase, suppress aberrant RAD51 recruitment, and prevent the accumulation of toxic HR intermediates that manifested as BLM-positive anaphase bridges^28^. We showed that AKT activity was required for 53BP1 NBs regulation, suppressing inappropriate RAD51 activity, and maintaining mitotic integrity. However, how p-AKT mechanistically controls 53BP1 NB function remains an important question for future study.

Altogether, we uncovered a novel role for nuclear F-actin and Anillin in the replication stress response: reshaping PML NB architecture, enabling AKT activation, and ensuring the faithful management of under-replicated DNA. More broadly, our work first established how direct physical interactions between actin filaments and nuclear condensates contributed to genome protection under replication stress.

## Acknowledgements

We thank the CMRI ACRF Telomere Analysis Centre supported by the Australian Cancer Research Foundation for microscopy infrastructure and support; the Westmead Institute of Medical Research Flow Cytometry Centre, supported by the Cancer Council NSW and the Australian NHMRC for cell sorting; the CMRI Biomedical Proteomics for mass-spectrometry and their assistance; CMRI Bioinformatics facility for assistance with data analysis; CMRI Vectorology Facility for manufacturing AAV viruses; Zdenek Lansky and Ravid Straussman for sharing plasmids. Figures 5j and Supplementary 9f were created with BioRender. Members of the Lamm and Cesare labs are thanked for their suggestions and discussion. N.L. is supported by the Col Reynolds Fellowship by The Kids’ Cancer Project. This project was supported by grants from the Hill Foundation for N.L.

## Author contributions

V.D.L.P., H.W.L. and N.L. conceived the study. V.D.L.P., H.W.L., B.J.E.L. and V.J.W. performed most of this manuscript’s experiments and data analysis. C.W. and D.G. performed all *in vitro* droplet assays. J.G. performed endogenous PML Co-IP. M.E.G. performed mass spectrometry experiments. L.L.M. and C.B.N. completed the mass spectrometry analysis. S.V., M.S., L.F. and S.R. created and verified plasmids and cell lines and assisted with minor experimentation. J.S.L. purified Anillin. J.S.L., M.B., H.A.P., J.C.S., A.J.C. and N.L. supervised junior members of the research team and secured funding. C.W., J.C.S., A.J.C. and N.L. wrote the manuscript.

## Methodes

### Cell culture and treatment

U-2OS, HeLa, U-2OS-Flag-APEX2-PML, U-2OS-Flag-APEX2, HeLa-Flag-APEX2-PML, HeLa-Flag-APEX2, U-2OS-miRFP-PML, HeLa-miRFP-PML, U-2OS-mStrawberry-FoxO1, IMR90 and SaOS2, cultures were grown in Dulbecco’s Modified Eagle Medium (DMEM) supplemented with 10% fetal calf serum (F9423, Sigma), GlutaMax and MEM non-essential amino acid solution (Gibco) and maintained at 37 °C in 10% CO_2_ and 3% O_2_. Cell Bank Australia verified cell line identity using short-tandem-repeat profiling, and cells were identified to be mycoplasma negative (MycoAlert, LT07-118, Lonza). Dimethyl sulfoxide (DMSO; Sigma-Aldrich), Aphidicolin (APH; Sigma-Aldrich), Hydroxyurea (HU; Sigma-Aldrich), Latrunculin A (LatA; Cayman Chemical, 10010631), VE-822 (Selleckchem, S7102), CK-666 (Sigma-Aldrich, SML0006), Ku-55933 (Sigma-Aldrich), and MK-2206 (Cayman Chemical, 11593) were used in cell treatments.

### Plasmid cloning

Plasmids used in this study are listed in Supplementary Table 1. The pFRT-lacZeo, pOG44, pcDNA6-TR, and pcDNA5/FRT/TO vectors were obtained from the Flp-In™ T-REx™ system (Thermo Fisher Scientific, K650001). FLAG-APEX2 was PCR-amplified from pRRL-GW-APEX2-TRF1^80^ and inserted into NotI-digested pcDNA5-FRT-EV to generate pFRT-T0– FLAG-APEX2. For pFRT-T0-miRFP670-PML, miRFP670 and PML fragments were amplified from pCMV Actin(R62D)-NLS-P2A-miRFP670^3^ and pCMV PML-myc-FLAG (OriGene, RC214606), respectively, and co-inserted into NotI-digested pcDNA5-FRT-EV. pLXSN-FLAG-APEX2 and pLXSN-FLAG-APEX2-PML were generated by PCR-amplifying the FLAG-APEX2 fragment from pFRT-T0-FLAG-APEX2^80^ and the FLAG-APEX2-PML fragment from pFRT-T0-FLAG-APEX2-PML and subcloning them into EcoRI-digested pLXSN (Takara, 631509), respectively.

To generate the plasmid pJSL096 encoding Anillin-GFP, the Anillin-GFP sequence was amplified from the pDL150 plasmid (a generous gift from Zdenek Lansky^60^). The amplified product was then cloned into the pGB-01;02-ccdB (a generous gift from Peggy Stolt-Bergner^81^, LMBP 12025, BCCM/GeneCorner).

All cloning was performed using In-Fusion cloning (Takara Bio) according to the manufacturer’s instructions, and multi-fragment constructs were sequence-verified to confirm insertion and correct orientation.

### Viral transduction and cell line generation

U2OS FRT-T0 and derivative cell lines created using the Flp-In™ T-REx™ system (Thermo Fisher Scientific, K650001). U2OS cells were transiently transfected with pFRT-lacZeo and selected in Zeocin (Gibco, Thermo Fisher Scientific, R25001; 100 µg/mL) to integrate one or more Flp recombination target (FRT) sites into the genome. Cells were subsequently transfected with pcDNA6/TR and selected in blasticidin (Sigma-Aldrich, Merck, SBR00022; 5 µg/mL) to integrate the tetracycline repressor (TetR) element stably, enabling doxycycline/tetracycline-inducible expression. The resulting parental line (U2OS FRT-T0) contained genomic FRT site(s) and a functional TetR element. For site-specific integration of the gene of interest (FLAG-miRFP670-PML, FLAG-APEX2, or FLAG-APEX2-PML), U2OS FRT-T0 cells were co-transfected with 8 µg total DNA at a 9:1 mass ratio of pOG44 (encoding Flp recombinase) to donor plasmid. Flp-mediated recombination between genomic and plasmid FRT sites yielded stably integrated transgenes. Hygromycin B (Roche, Merck, 10843555001; 200 µg/mL) was used to select for stable integrants. Resistant cell populations were pooled and expanded, and expression was induced with 1 µg/mL doxycycline (Sigma-Aldrich, Merck, D3072) for 48 h.

We were unsuccessful in establishing the FRT-T0 platform in HeLa cells and therefore used an alternative strategy. Parental HeLa cells were transduced with retroviruses encoding pLXSN-FLAG-APEX2, pLXSN-FLAG-APEX2-PML, or pLXSN-RFP-PML. Retroviral particles were generated by transfecting Phoenix packaging cells in T75 flasks with 20 µg of the corresponding plasmid using Lipofectamine 3000 (Invitrogen, Thermo Fisher Scientific). Viral supernatants were collected, passed through a 0.45 µm syringe filter, and applied to HeLa cells seeded in 6-well plates in the presence of 8 ug/mL polybrene (Sigma-Aldrich, Merck, H9268) at a 1:1 ratio with DMEM. Transduced cells were selected with Geneticin (G418; Roche, Merck, A1720; 400 µg/mL), and surviving colonies were pooled to establish stable cell lines.

The AKT translocation reporter U-2OS cell line was made by transducing parental U2OS cells with lentivirus carrying the pLKO.1-FoxO1-mStrawberry-pGK-BSD construct (a generous gift from Ravid Straussman^76^; Addgene, Plasmid #158685) and selecting in 5 ug/mL Blasticidin.

The 3F U-2OS cells were created using tFucci(CA)2/pCSII-EF^48^ provided by the RIKEN BRC through the National BioResource Project of the MEXT, Japan (cat. RDB15446). The third-generation lentivectors harbouring tFucci(CA)2/pCSII-EF were produced in the CMRI Vector and Genome Engineering Facility. The cultures were transduced for 48 h with crude supernatants in media supplemented with 4 µg ml^−1^ of polybrene. Cells were cultured for 6 days and sorted at the Westmead Institute for Medical Research Flow Cytometry Facility for mCherry expression. Appropriate cell cycle-dependent colourimetric signalling verified by live imaging.

### siRNA transfection

A transient gene knockdown was achieved using non-targeting (control siRNA, D-001810-10), *PML*(L-006547-00-0010), *Anillin* (L-006838-00-0010), *AKT1* (L-003000-00-0010), *AKT2* (L-003001-00-0010) and *AKT3* (L-003002-00-0010) ON-TARGETplus siRNA pools (Dharmacon) at the final concentration of 10 nM. Cells were transfected using Lipofectamine RNAiMAX (Invitrogen Thermo Fisher Scientific, 13778150) according to the manufacturer’s instructions.

### Cell cycle synchronisation

The cells were synchronised at G1/S using a double thymidine block by treating with 2 mM thymidine for 16-18 h, releasing for 6-8 h and treating again with 2 mM thymidine for 16-18 h.

### Phalloidin staining

Cells were seeded on sterile glass coverslips, treated with APH 24 h later, and then fixed 24 h thereafter. Cells were fixed for 1 min in cytoskeleton buffer (10 mM MES, 150 mM NaCl, 5 mM EGTA, 5 mM glucose and 5 mM MgCl_2_, pH 6.1) plus 0.5% Triton X-100 and 0.25% glutaraldehyde (Sigma-Aldrich), and then for 15 min in cytoskeleton buffer plus 2% glutaraldehyde. Autofluorescence was quenched with fresh 1 mg ml^−1^ sodium borohydride (Sigma-Aldrich) and cover slips were incubated overnight at 4 °C with phalloidin-Atto 488 (Sigma-Aldrich, 49409). Slides were washed four times for 5 min in 1× PBS and incubated for 3 h at 37 °C with anti-lamin A/C (Sigma-Aldrich SAB4200236) and PML (Santa-Cruz sc-966) antibodies diluted in 1× PBS plus 5% fetal calf serum (1:200). The samples were washed again in the same manner and then incubated with Alexa Fluor 450 and 568-conjugated secondary antibodies for 1 h at 37 °C. Slides were washed as described above, dehydrated in a graded ethanol series (70%, 90% and 100% for 2 min in each solution) and mounted with Prolong Gold (Life Technologies). To avoid complications of extranuclear signals, imaging was limited to cells with a clearly resolvable nuclear interior.

### Chromobody transfection and imaging

The NLS-GFP-actin chromobody (Chromotek, acg-n, nuclear-actin-CB) and RFP-PCNA chromobody (Chromotek, ccr, PCNA-CB) were purchased with a material transfer agreement from Chromotek. Plasmids were transfected with Lipofectamine LTX (Thermo Fisher Scientific). For fixed imaging, cells were seeded on sterile cover slips and transfected 24 h later. Chemical inhibitors were added 24 h after transfection for 24 h. Samples were fixed in 3.7% formaldehyde/1× PBS for 10 min, permeabilised with 0.5% Triton X-100/1× PBS for 10 min at room temperature and blocked with 5% bovine serum albumin (Sigma-Aldrich)/1× PBS. Samples were incubated with primary PML (Santa-Cruz sc-966) or p-AKT-Ser473 (Cell Signalling 9271) antibodies (1:200) and secondary antibodies (1:500) before dehydrated in a graded ethanol series (70%, 90% and 100%) and then mounted using Prolong Gold. For live-cell imaging, cells were seeded on a glass-bottom dish (3 cm; World Precision Instruments) and transfected 24 h later. Then, 48 h after transfection, the culture medium was replaced with colourless DMEM (FluoroBrite DMEM, Life Technologies) supplemented with the indicated compounds. Imaging was performed at 37 °C, 10% CO_2_ and 3% O_2_ on a Zeiss Cell Observer SD spinning-disk confocal microscope using combined differential interference contrast and fluorescence imaging (a 561 nm laser, 7% excitation power, 1 × 1 binning, EM gain of 908; and a 488 nm laser, 6.5% excitation power, 1 × 1 binning, EM gain of 527) with appropriate filter sets and a ×63/1.3 NA oil-immersion objective. Ten images per *z* stack were captured in an image scaled to 47,504 px × 37,602 px at 10.05 μm × 7.96 μm using Zen Blue 2 v.2.0.14283.302 (ZEISS) and an Evolve Delta (Photometrics) camera every 90–200 s for up to 72 h. F-actin^+^ nuclei represent the fraction of S-phase nuclei or G1-phase nuclei that were actin-fibre positive at least once during imaging. For G1 nuclei

### PML, p-AKT, 53BP1 and γ-H2AX labelling

24 h after APH addition, cells grown on sterile glass cover slips were fixed in 3.7% formaldehyde/1× PBS for 10 min, permeabilised with 0.5% Triton X-100/1× PBS and incubated with one or two of the following primary antibodies: PML (Santa-Cruz sc-966), p-AKT-Ser473 (Cell Signalling 9271), p-AKT-Ser473 (Proteintech, 28731-1-A), 53BP1 (Thermo Scientific, MA5-26830), γ-H2AX (abcam, ab2893). Cells were then incubated with secondary antibodies, washed, dehydrated and mounted as described above.

### Super-resolution imaging

Super-resolution imaging was performed on a ZEISS LSM 880 AxioObserver confocal fluorescent microscope fitted with an Airyscan detector using a Plan-Apochromat ×63/1.4 NA M27 oil-immersion objective using ZEN Black 2.3 pro SP1 v.14.0.20.201 (ZEISS). Cells were imaged using 1.9% excitation power of 568 nm laser, 2% excitation power of 488 nm laser and 1.8% excitation power of 647 nm laser, with 1 × 1 binning for all laser conditions in combination with the appropriate filter sets. Ten or more z stacks were captured with frame scanning mode and unidirectional scanning. The z stacks were Airyscan processed using batch mode in Zen software.

### Quantitative Image-Based Cytometry (QIBC)

Cells grown on sterile glass coverslips were treated with DMSO or APH, LatA, CK666 or combined for 24 h before fixation. Alternatively, cells were transfected with the appropriate siRNA 72 h before fixation. In the last hour before fixation, cells were pulse-labelled with 10 µM EdU. Cells were then fixed immediately (4% paraformaldehyde in PBS for 10 min at 4 °C). All the wash steps were performed in PBS with 0.1% Tween. The samples were washed three times, permeabilised with 0.5% Triton X-100 in PBS + 0.1% Tween (PBST) for 5 min, rewashed three times and blocked with 2% BSA buffer (in PBST) for 2 h. The samples were incubated overnight at 4 °C with one of the following primary antibodies: PML (Santa-Cruz sc-966), p-AKT-Ser473 (Cell Signalling 9271), 53BP1 (Thermo Scientific, MA5-26830), RAD51 antibody (Calbiochem, PC130, 1:500 in 2% BSA), then washed five times. All subsequent steps were performed in the dark. The coverslips were incubated with a click-iT EdU solution of 10 μM 6-carboxyfluorescein-TEG azide (Berry and Associates, FF 6110), 10 mM sodium ʟ-ascorbate (Sigma-Aldrich, A4034), 2 mM copper (II) sulphate (Sigma-Aldrich, 451657 in PBS) for 30 min. The cells were washed five times in 1% BSA (in PBST) before staining with Alexa Fluor 488-conjugated secondary antibody (Invitrogen, A11036, 1:500 in 2% BSA) for 1.5 h. The cells were washed three times, incubated with 4,6-diamidino-2-phenylindole (DAPI; Sigma-Aldrich, 10236276001) for 5 min (1:5000 in PBST), washed 1×, rinsed twice with milliQ H_2_O and sequentially dehydrated in 70%, 90% and 100% ethanol for 3 min each. The air-dried coverslips were mounted with Prolong Gold Antifade (Invitrogen, Thermo Fisher Scientific, P36934) and cured in the dark for ≥24 h. The fixed images were acquired with a ZEISS AxioImager Z.2 microscope fitted with a 40× 1.3 NA oil objective, Axiocam 506 monochromatic camera using ZEN Blue v2.3 pro software. A total of 19 *Z* stacks for each channel were taken at 0.24 µm increments.

The extended depth of focus function in ZEN Blue v2.3 pro software was used to make maximum intensity projections of each image, and then CZI files were converted to single-channel. TIFF files and imported into CellProfiler v4.2.1. Using custom image analysis pipelines, we pre-processed images using the CorrectIllumination and EnhanceOrSuppress feature functions. The nuclei and subnuclear foci were identified using the IdentifyPrimaryObjects functions with either Minimum Cross-Entropy or Otsu thresholding. The foci were associated with their respective nuclei using the Relate function, and the foci intensities were measured using the MeasureObjectIntensity function. The data were plotted in GraphPad Prism v10.6.0 using a double-gradient colourmap for the multiple variables plots.

Cell cycle analysis was not conducted in si-ANLN experiments, as the multinucleated phenotype confounds accurate assessment.

### Visualising BLM positive anaphase bridges

Cells were transfected with siRNA, synchronised by a double-thymidine block, and seeded onto sterile glass coverslips. Following synchronisation, cells were treated with either DMSO or APH for 22 h, then fixed, or released into drug-free medium for 24 h before fixation. Cells were incubated with BLM antibody (Abcam, ab2179), followed by the appropriate secondary antibody, then washed, dehydrated, and mounted as described above.

Samples were imaged using a ZEISS AxioImager Z.2 with a ×63/1.4 NA oil-immersion objective, appropriate filter cubes, an Axiocam 506 monochromatic camera and Zen 2.3 Pro v.2.3.69.01015. Images were scored by eye.

### Preparation of nuclear and cytoplasmic extracts

24 hours after drug treatment or 72 h following siRNA transfection, cells were harvested by trypsinisation, pelleted by centrifugation, counted with a Countess automated cell counter, and washed once in PBS. Pellets were resuspended in 20 volumes of hypotonic buffer (10 mM HEPES-KOH, pH 8.0, 10 mM KCl, 1 mM MgCl₂, 1 mM DTT; Invitrogen P2325) and incubated on ice for 10 min to allow swelling. Plasma membranes were lysed by adding one-tenth volume of 5% Triton X-100, and nuclei were pelleted at 1,000 g for 1 min. Approximately 75% of the supernatant was collected as the cytoplasmic fraction, and the remainder was discarded.

Nuclear pellets were washed 2-3 times with hypotonic buffer lacking Triton X-100 and repelleted at 1,000 g for 1 min, with supernatants carefully aspirated to avoid nuclear loss. Both nuclear and cytoplasmic fractions were snap-frozen and stored at –80 °C. For downstream analysis, fractions were solubilised in 4× LDS (Laemmli) buffer supplemented with DTT (1:20 dilution from a 1 M stock) and Benzonase (1:100 dilution; Merck E1014, 25 KU). Nuclear fractions were eluted in 50 µL (corresponding to one-quarter of the initial resuspension volume). Cytoplasmic fractions were prepared by mixing 50 µL lysate with 50 µL 4× LDS buffer, and twice as much cytoplasmic material was loaded relative to nuclear samples for western blotting, yielding a nuclear-to-cytoplasmic loading ratio of 4:1.

### Western blotting

The whole-cell extract preparation and western blots were performed as described previously^82^. Luminescence signals were visualised on an LAS 4000 imager (Fujifilm). Primary antibodies include: Anillin (E9L6E, Cell Signaling Technology, 85785, 1:1,000), PML (EPR16792, Abcam 179466, 1:1,000), DAXX (Sigma-Aldrich, HPA00873, 1:1,000), AKT (Cell Signaling Technology, 9272, 1:1,000), p-AKT-S473 (Cell Signaling Technology, 9271, 1:1,000), β-actin (AC-15, Sigma-Aldrich, A5441, 1:10,000), Vinculin (Sigma-Aldrich, V9131, 1:10,000), CHK1 (2G1D5, Cell Signaling Technology, 2360, 1:1,000), p-CHK1-pS345 (133D3, Cell Signaling Technology, 2348, 1:1,000), CHK2 (Cell Signaling Technology, 2662, 1:1000), CHK2-pT68 (Cell Signaling Technology, 2661, 1:1000) cyclin A2 (EPR17351, Abcam, ab181591, 1:3,000), GAPDH (D16H11, Cell Signaling Technology, 5174, 1:5,000), histone H2B (V119, Cell Signaling Technology, 8135, 1:1,000),, The secondary antibodies include horseradish peroxidase conjugated goat anti-mouse (Dako, P0447, 1:5,000–1:20,000) and goat anti-rabbit (Dako, P0448, 1:5,000–1:20,000).

### PML co-immunoprecipitation

Cells were treated with DMSO, APH or a combination of APH with CK666 for 24 hours before equal numbers of cells for each treatment were harvested and snap frozen. The cells were lysed using lysis buffer B (20mM of HEPES-KOH (pH 7.9), 200mM NaCl, 2mM MgCl2, 10% v/v glycerol, 0.1% Triton X-100, 1mM dithiothreitol, cOmplete™, EDTA-free Protease Inhibitor cocktail (Roche, catalogue # 04693132001)) for 1 hour at 4 °C with vortexing. Following pre-clearing by centrifugation for 40 minutes at 4 °C and 13000rpm, the lysis supernatant was collected as input and for co-immunoprecipitation. Pull-down of PML proteins was performed using protein G Dynabeads (Invitrogen, catalogue # 10004D). For each sample, 50 µl of the bead slurry was washed twice in PBST before 3 µg of anti-PML antibody (Abcam, Ab179466) or rabbit IgG isotype control (Cell Signaling Technology, 2729S) was added to the beads in 200 µl of PBST and incubated for 30 min at room temperature with rotation. Beads were then washed 3x with PBST before the antibodies were crosslinked onto the beads using Bis(sulfosuccinimidyl)suberate (BS3) (Thermo Fisher Scientific, 21580). Specifically, BS3 was dissolved in conjugation buffer (20mM sodium phosphate, 150mM NaCl, pH 7-9) to 5mM, fresh on the day of use. Beads were washed twice with conjugation buffer before incubation in 240µl of 5mM BS3 for 30 minutes at room temperature with rotation. The crosslinking reaction was quenched by incubation with 80 µl of 1M tris pH 7.5 at room temperature for 30mins. The beads were then washed 3-5 times with PBST before the protein lysate was incubated with the beads overnight, at 4 °C with rotation. The next day, the beads were washed 3x with cold lysis buffer B before being heated to 75 °C for 10 minutes in 1x NuPage elution buffer (Thermo Fisher Scientific, NP0007) to elute the bound proteins. Western blots were then run on elution products and inputs.

### Anillin protein purification

To purify Anillin-GFP, we used the GoldenBac baculovirus expression system^81^ in Tni cells (expression systems). Cells were infected at a density of 1×10^6^ cells per mL and incubated for 72 hours post-infection. Subsequently, the cell pellet from a 1 L culture was resuspended in 50 mL of ice-cold His-Trap buffer (50 mM Tris-HCl, pH 8.0; 5% glycerol; 300 mM NaCl; 1 mM MgCl_2_; 0.005% Tween-20) supplemented with 20 mM imidazole, a protease inhibitor cocktail (cOmplete, EDTA-free, Roche), and benzonase at a final concentration of 25 units/mL and lysed by dounce homogenisation. The lysate was centrifuged at 80,000×g for 30 minutes at 4°C to obtain a clarified supernatant. The supernatant was applied to a 5 mL Ni-NTA column (Cytiva) equilibrated with His-Trap buffer containing 20 mM imidazole. After washing the column with 10 column volumes (CV) of His-Trap buffer, bound proteins were eluted using His-Trap buffer supplemented with 300 mM imidazole. Fractions containing Anillin-GFP were pooled and diluted to achieve a final NaCl concentration of 50 mM. The diluted protein solution was loaded onto a 1 mL Hi-Trap Q anion exchange column (Cytiva) equilibrated with buffer containing 50 mM Tris-HCl (pH 8.0), 100 mM NaCl, 10% glycerol, 0.5 mM TCEP, and 0.01% NP-40-S. The column was washed with 10 CV of the same buffer, and Anillin-GFP was eluted using a linear gradient from 100 mM to 1,000 mM NaCl over 30 CV. Peak fractions were pooled and concentrated to approximately 2 mL using an Amicon Ultra 30 kDa MWCO concentrator. The concentrated protein was dialysed overnight into storage buffer comprising 50 mM Tris-HCl (pH 8.0), 300 mM NaCl, 0.5 mM EDTA, 1 mM MgCl_2_, 1 mM DTT, and 5% glycerol. Protein concentration was determined using a NanoDrop spectrophotometer.

### APEX2-PML proximity biotin-labelling

U2OS FRT-T0 FLAG-APEX2-PML and U2OS FRT-T0 FLAG-APEX2-EV cells were pre-treated with doxycycline (1 µg/mL) for 48 h to induce expression, whereas HeLa FLAG-APEX2-PML and HeLa FLAG-APEX2-EV cells were not inducible and therefore cultured without doxycycline. Cells were then incubated in biotin-phenol (Iris Biotech, LS-3500; 500 μM)-supplemented medium for 30 min at 37 °C, and labelling was initiated by the addition of freshly prepared hydrogen peroxide (1 mM final) for 1 min with gentle swirling. The reaction was quenched by the addition of 2× quench buffer containing 20 mM sodium ascorbate (Sigma-Aldrich, Merck, A4034), 10 mM Trolox (Sigma-Aldrich, Merck, 238813), and 20 mM sodium azide (Sigma-Aldrich, Merck, S2002), followed by gentle mixing for 30 s. After aspiration, cells were washed twice for 2 min in 1× quench buffer (10 mM sodium ascorbate, 5 mM Trolox, 10 mM sodium azide), rinsed with PBS, and fixed for immunofluorescence, harvested for protein extraction for western blot or LC-MS/MS.

### Recovery of biotinylated proteins

After labelling, cells were lysed directly on the plate in 2 mL of SDC buffer (4% sodium deoxycholate, 100 mM Tris-HCl pH 8.5), collected by scraping, and transferred into 2 mL tubes. Lysates were mixed by pipetting, heated at 95 °C for 5 min, cooled on ice, and this heat-cool cycle was repeated once before storage at −80 °C. For proteomic preparation, lysates were thawed on ice and homogenised using a Bioruptor sonicator for 20 min at maximum output, performed in batches to accommodate sample capacity. Protein concentration was determined using a BCA assay (Pierce™ BCA Protein Assay Kit, Thermo Scientific, Thermo Fisher Scientific, 23225), and 1 mg of total protein from each sample was aliquoted and adjusted to 270 µL in SDC lysis buffer. Reduction and alkylation were performed by adding 10% sample volume of freshly prepared buffer containing 100 mM tris(2-carboxyethyl)phosphine hydrochloride (TCEP-HCl; Thermo Fisher, Thermo Fisher Scientific, 20490), 400 mM 2-chloroacetamide (CAA; Sigma-Aldrich, Merck, C0267), and 400 mM KOH, followed by incubation at 45 °C for 5 min with shaking at 1500 rpm. Streptavidin magnetic beads (Pierce™ Streptavidin Magnetic Beads, Thermo Scientific, Thermo Fisher Scientific, 88817) were washed in SDC lysis buffer, divided into aliquots, and incubated with the reduced and alkylated lysates overnight at 4 °C with shaking at 1200 rpm. The following day, beads were washed three times with SDC lysis buffer and maintained on ice prior to downstream processing (western blots or mass spectrometry).

### Sample preparation and peptide labelling for mass spectrometry

Samples were washed in 500 µL of 50 mM 4-(2-hydroxyethyl)-1-piperazineethanesulfonic acid (HEPES). To each sample, 40 µL of 50 mM HEPES containing 10 mM tris (2-carboxyethyl)phosphine (TCEP) was added, and samples were heated at 85 °C for 10 min. After cooling, iodoacetamide was added to a final concentration of 20 mM, and samples were incubated at 23 °C for 30 min in the dark. Digestion was performed by adding 1 µg of trypsin (Merck, Trypzean) and incubating at 37 °C for 3 h, followed by an additional 1 µg of trypsin and incubation for 8 h at 37 °C.

Peptides were collected, and an additional 40 µL of 50 mM HEPES was added to recover residual peptides. Samples were then labelled with TMTpro reagents (Thermo Fisher Scientific) according to the manufacturer’s protocol. After confirming efficient labelling, samples were combined, desalted using solid-phase extraction (Oasis PRIME HLB, 1 cc, 30 mg; Waters), and subjected to hydrophilic interaction liquid chromatography (HILIC) fractionation.

Fractionation was performed on a Dionex Ultimate 3000 HPLC system using a TSKgel Amide-80 column (250 mm × 1 mm, Tosoh Biosciences). Buffer A consisted of 90% acetonitrile with 0.1% trifluoroacetic acid (TFA), and Buffer B was 0.1% TFA in water. Samples were loaded in Buffer A at a flow rate of 60 µL/min for 10 min. The gradient was run from 100% Buffer A to 60% Buffer A over 35 min at 50 µL/min. Fractions were collected into a 96-well plate at 30 s intervals using a Probot fraction collector (LC Packings) and monitored by UV absorbance at 214 nm. A total of 27 fractions were obtained, which were then combined into pools of approximately equal peptide content based on UV signal. Fractions were dried and reconstituted in 0.1% formic acid prior to LC–MS/MS analysis.

### LC-MS/MS

The LC-MS/MS was performed using a Dionex UltiMate 3000 RSLC nano system and Q Exactive Plus hybrid quadrupole-orbitrap mass spectrometer (Thermo Fisher Scientific). Each HILIC fraction was loaded directly onto an in-house 300 mm long 0.075 mm inside diameter column packed with ReproSil Pur C18 AQ 1.9 μm resin (Dr Maisch, Germany). The column was heated to 50 °C using a column oven (PRSO-V1, Sonation lab solutions, Germany) integrated with the nano flex ion source with an electrospray operating at 2.3 kV. The S lens radio frequency level was 50, and the capillary temperature was 250 °C. The sample was loaded onto the column in 99% reversed-phase buffer A (solution of 0.1% formic acid) and 1% buffer B (solution of 0.1% formic acid, 90% acetonitrile). The loading was for 17.5 min at 300 nL/min using 1% buffer B. The gradient, at 250 nL/min, was from 1% buffer B to 7% buffer B in 1 min, to 29% buffer B in 101.5 min, to 36% buffer B in 8 min, to 99% buffer B in 1 min, held at 99% buffer B for 2 min, to 99% buffer A in 1 min and held for 8 min as the flow rate increased to 300 nL/min. MS acquisition was performed for 140 min. All samples and fractions were analysed using data-dependent acquisition LC-MS/MS. The MS scans were at a resolution of 70,000 with an automatic gain control target of 1,000,000 for a maximum ion time of 100 ms from m/z 375 to 1500. The MS/MS scans were at a resolution of 35,000 with an automatic gain control target of 200,000 and maximum ion time of 100 ms. The loop count was 12, the isolation window was 1.1 m/z, the first mass was fixed at m/z 120, and the normalised collision energy was 31. Singly charged ions and those with charge >8 were excluded from MS/MS and dynamic exclusion was for 37 s.

The raw LC-MS/MS data were processed with MaxQuant v1.6.7.0. Variable modifications were oxidation (M), acetyl (protein N-terminus) and deamidation (NQ). Carbamidomethyl (C) was a fixed modification. Digestion was set to trypsin/P with a maximum of 3 missed cleavages. The TMTpro correction factors were entered (lot VJ313476 and XC343801). Minimum reporter peptide ion fraction was 0.6. The *Homo sapiens* reference proteome with canonical and isoform sequences downloaded Aug 21, 2023, with 82,427 entries and 20,586 genes. The inbuilt contaminants fasta file was also used. A second peptide search was enabled. The peptide spectrum matching and protein false discovery rates were 1%. All modified peptides and counterpart non-modified peptides were excluded from protein quantification. All other settings were default within MaxQuant v1.6.7.0.

### Proteomic preprocessing and normalisation

Proteomic analysis was performed in the Bioinformatics Core Facility at the Children’s Medical Research Institute. Protein reporter intensity corrected values from MaxQuant were imported into R (v4.4.3) for downstream analysis. Proteins identified by only modified peptides, reverse peptides or potential contaminant peptides were removed. Proteins with unique peptide counts below 1 were also discarded for the analysis. For multimapped proteins, the UniprotKB protein accessions were kept if they contained a unique gene symbol. If duplicated gene symbols existed, a single UniprotKB protein accession was maintained based on the highest protein evidence, reviewed status, canonical status and then the lowest isoform number. Proteins were filtered for any row containing more than zero missing values. Proteomic intensities were log2 transformed and Vsn normalised using the limma package (v3.62.1). An Anova model was used to identify 500 negative control genes (Adjusted p-value >= 0.5). These negative control genes were then used to run RUVIII unwanted variation removal, adjusting for wanted components (k = 3 for U-2OS and k = 4 for HeLa) from the ruv package (v0.9.7.1).

### Proteomic differential expression analysis

Differential protein abundance between experimental groups was assessed using the limma package. The log2-transformed proteomic intensities were fitted to a linear model, with empirical Bayes moderation to shrink variance estimates across proteins, and adjusted p-values were computed using the Benjamini-Hochberg method to control for false discovery rate. Significant differentially expressed proteins were defined as those with adjusted p-values less than 0.05 and log2 fold changes less than negative 0.5 or greater than 0.5 (Adjusted p-value < 0.05 and -0.5 < Log2 Fold Change > 0.05).

### Proteomic odds ratio analysis

Significantly regulated proteins (Adjusted p-value < 0.05 and -0.5 < Log2 Fold Change > 0.05) were subjected to an odds ratio analysis to identify enriched pathways using the clusterProfiler package (v4.14.0). Multimapped proteins were separated into individual proteins, and for each gene symbol, the most significant (lowest adjusted p-value) was used for the pathway analysis. Pathway enrichment was performed separately for both significantly downregulated and upregulated proteins, against the GO, KEGG and REACTOME databases, to identify significantly enriched pathways (Adjusted p-value < 0.05).

### Proteomic gene set enrichment analysis

Gene Set Enrichment Analysis was used to identify enriched pathways using the clusterProfiler package (v4.14.0). Multimapped proteins were separated into individual proteins, and for each gene symbol, the most significant (lowest adjusted p-value) was used for the pathway analysis. Proteins were then ranked by the sign of the log2 fold change times by negative log10 p-value. This ranked protein list was then used for pathway enrichment, against the GO, KEGG and REACTOME databases, to identify significantly enriched pathways (Adjusted p-value < 0.05).

### Actin Labelling

Monomeric actin (Rabbit muscle actin, Cytoskeleton) was labelled using maleimide-conjugated dyes (Sigma-Aldrich). Dyes were incubated with G-actin at a 2-fold molar excess for 2 hours at room temperature before being separated from the labelled protein by applying the labelling reaction to a spin column packed with Sephadex G-50 Fine DNA Grade (GE Healthcare GE17-0573-01) hydrated with A buffer (5 mM Tris-HCL (pH 8), 0.2 mM ATP and 0.5mM DTT (pH 8)). The labelled protein was then centrifuged at 100,000 x G for 10 min at 4 °C to remove aggregates before being flash-frozen in single-use aliquots.

### Protein condensate formation and actin filament assembly

Condensates composed of Anillin were formed by mixing the indicated concentration of protein (see text) with 3% (w/v) PEG 8000 in 20 mM Tris (pH 7.4), 5 mM TCEP, and the given concentration of NaCl. PEG was added last to induce condensate formation after the protein was evenly dispersed in the solution.

For actin assembly assays within condensates, condensates were formed for ten minutes (with time starting after PEG addition), and then G-actin was added to the condensate solution and allowed to assemble for 15 minutes before imaging. For phalloidin-actin assays, unlabelled G-actin was added to pre-formed protein condensates and allowed to assemble for 10 min. Phalloidin-iFluor594 (Sigma-Aldrich) was then added to stain filamentous actin for 10 min before imaging.

For FRAP experiments, A region within Anillin condensates was bleached, and consecutive images were taken every three seconds to monitor fluorescence recovery over time.

### Condensate imaging

Samples were prepared for microscopy in 3.5mm diameter wells formed using biopsy punches to create holes in 1.6 mm thick silicone gaskets (Grace Biolabs) on Hellmanex III cleaned, no. 1.5 glass coverslips (VWR). Coverslips were passivated using poly-L-lysine conjugated PEG chains (PLL-PEG). To prevent evaporation during imaging, an additional small coverslip was placed on top of the gasket to seal the well. Fluorescence microscopy was done using an Olympus SpinSR10 spinning disk confocal microscope with a Hamamatsu Orca Flash 4.0V3 Scientific CMOS camera. FRAP was done using the Olympus FRAP unit 405 nm laser.

PLL-PEG was prepared as described previously with minor alterations^56^. Briefly, amine-reactive mPEG succinimidyl valerate (Laysan Bio) was conjugated to poly-L-lysine at a molar ratio of 1:5 PEG to PLL. The conjugation reaction occurs in 50 mM sodium tetraborate solution, pH 8.5, and is allowed to react overnight at room temperature while continuously stirring. The final product is then buffer exchanged to PBS pH 7.4 using 7000 MWCO Zeba spin desalting columns (Thermo Fisher) and stored at 4 °C.

### Condensate image analysis

Image J was used to quantify the distribution of condensate characteristics. Specifically, condensates were selected using thresholding in the brightest channel and shape descriptors (i.e. diameter, aspect ratio, etc.), and protein fluorescent intensities were measured using the built-in analyse particles function. For aspect ratio analysis, condensates that had come into contact with other condensates were removed from the analysis to avoid any skewing of data from misrepresentation of single condensate deformation.

FRAP data were analysed using ImageJ, where fluorescence recovery over time was measured and then normalised to the maximum pre-bleach intensity. Recovery was measured for condensates of similar diameters and photobleached region size.

Partitioning data was calculated using the average intensities of the condensed protein phase and the bulk solution, with partitioning defined as the ratio of the intensity inside the condensate to outside the condensate. Images were cropped so that only condensates from the middle ninth of the field of view were analysed to avoid any error from potential non-uniform illumination across the imaging field.

### IncuCyte proliferation

Cells were imaged using an IncuCyte live imaging system (Sartorius) at 37 °C, 10% CO_2_ and 3% O_2_, with the images captured at 10× magnification every 2 or 4 h for the experiment duration. The proliferation was calculated using a confluence mask generated with the IncuCyte Zoom software (Sartorius, version 2019B). Inhibitors were added just prior to the start of imaging.

### Viability assay

Cells were seeded at 250 µL per well in black-walled, clear-bottom 96-well plates. The following day, cells were treated with combinations of MK-2206 (6 different concentrations) and hydroxyurea (HU; 9 different concentrations) and incubated for 72 h. Resazurin (Sigma-Aldrich) was then added to a final concentration of 10 µg/mL, and plates were incubated for 6 h before fluorescence measurement using EnSpire Multimode Plate Reader and EnSpire Manager v.4.13.3005.1482 (PerkinElmer; excitation 544 nm, emission 595 nm). IC₅₀ values were calculated using GraphPad Prism.

### Colony Formation Assays

Cells were seeded 1000 cells per well into a 6-well plate and treated with either DMSO, APH, MK2206 or combined. Wells were washed, supplemented with fresh media, and incubated until colonies were observed. Colonies were fixed with 4% PFA and stained with 0.05% Crystal violet.

### Statistics and reproducibility

All statistical analyses were performed using GraphPad Prism v10.6.0. Statistical methods and the number of cells (*N*), biological replicates (*n*) and error bars are described in the figure legends. The violin and box plots represent a cumulative distribution of data points from all replicates per condition. The representative data, whenever shown, are characteristic of similar results from the indicated number of independent biological replicates as described in the figure legends. The samples were randomly allocated into experimental groups. The data analysis was randomised as follows. In live imaging experiments, the cells within the field of view were selected randomly before analysis and followed for the experimental duration. To quantify PML NBs, p-AKT, 53BP1 and RAD51 foci, all interphase cells completely contained within the field of view were analysed using automated methods. All experiments were performed at least three times, unless specified otherwise, with quantification and statistics derived from *n* biological replicates or *N* cells, as described in the figure legends. For continuous data, parametric tests (*t*-test or analysis of variance (ANOVA) followed by Tukey’s post hoc test) were applied. The researchers were not blinded due to impracticability. The figures were prepared using Adobe Photoshop v25.0 and Illustrator v27.9.

