## Supplemental Figures for "Anillin-dependent actin assembly at PML NBs protects genome stability"

### SUPPLEMENTARY FIGURES AND FIGURE LEGENDS

#### Supplementary Figure 1

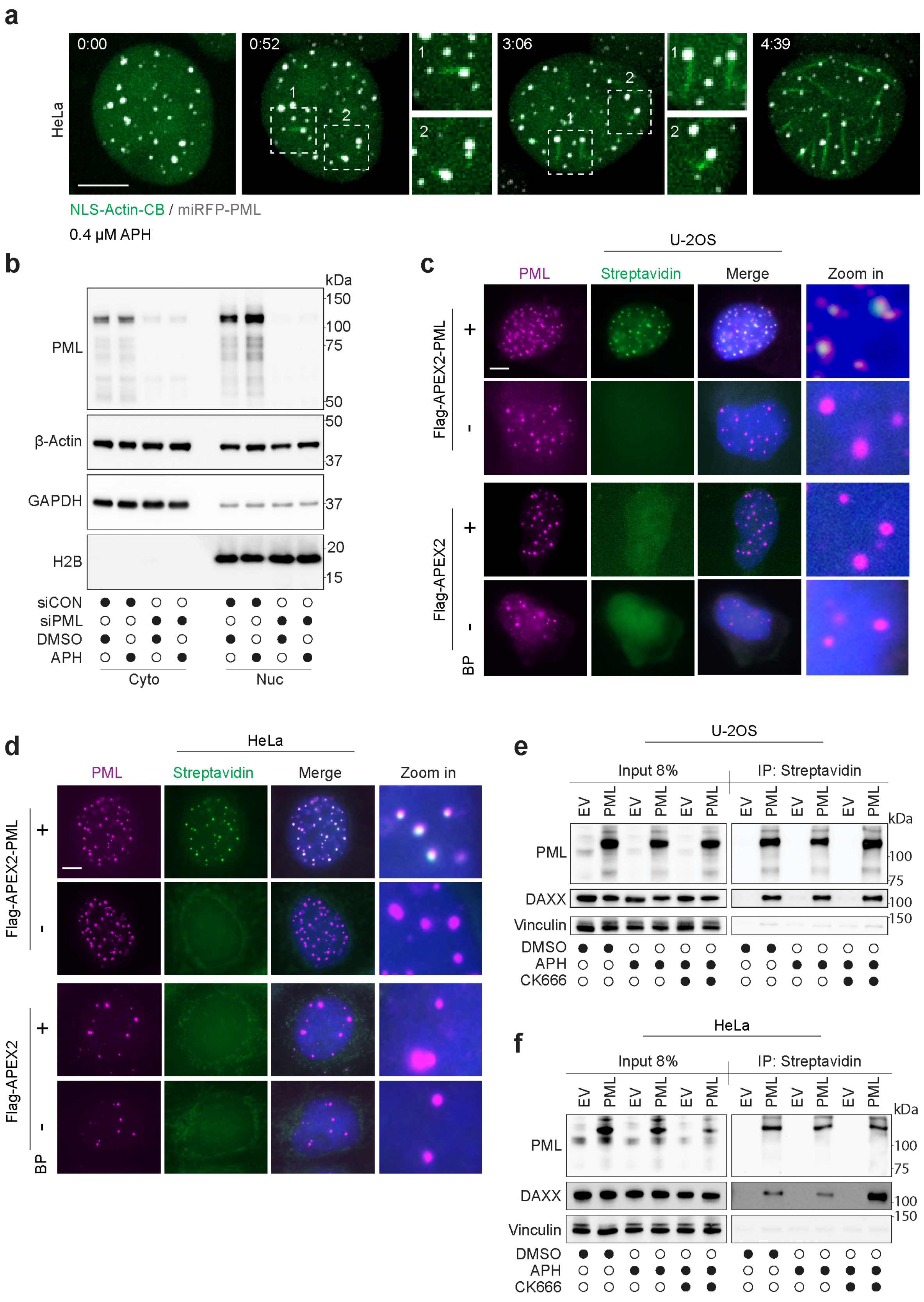

**Supplementary Figure 1 | Validation of stably expressed Flag-APEX2-PML U-2OS and HeLa cell lines.**

**a,** Representative live-cell spinning disk confocal microscopy images of HeLa cells expressing NLS-actin-CB and miRFP670-tagged PML treated with 0.4 µM APH for 24 h. Time is indicated in hours:minutes relative to the first time point. Scale bar 5µm.

**b**, Representative immunoblots of nuclear and cytosolic fractions from U-2OS cells transfected with si-PML or si-Control 72 h before extraction and treated with or without 0.4 µM aphidicolin (APH) for 24 h. The nuclear-to-cytoplasmic extract volume ratio loaded was 4:1, n = 3 biological replicates.

**c, d,** Immunofluorescence micrographs of U-2OS (**c**) and HeLa (**d**) cells stably expressing Flag-APEX2 or Flag-APEX2-PML, stained for PML and streptavidin (SA) with or without APEX activation by biotin-phenol (BP), n = 3 biological replicates.

**e, f,** Representative immunoblots of biotin-streptavidin immunoprecipitates from U-2OS (**e**) and HeLa (**f**) cells expressing Flag-APEX2 (Empty Vector; EV) or Flag-APEX2-PML (PML), treated for 24 h with DMSO, 0.4 µM APH, or APH in combination with 200 µM CK-666, n = 3 biological replicates.

**Supplementary Figure 2**

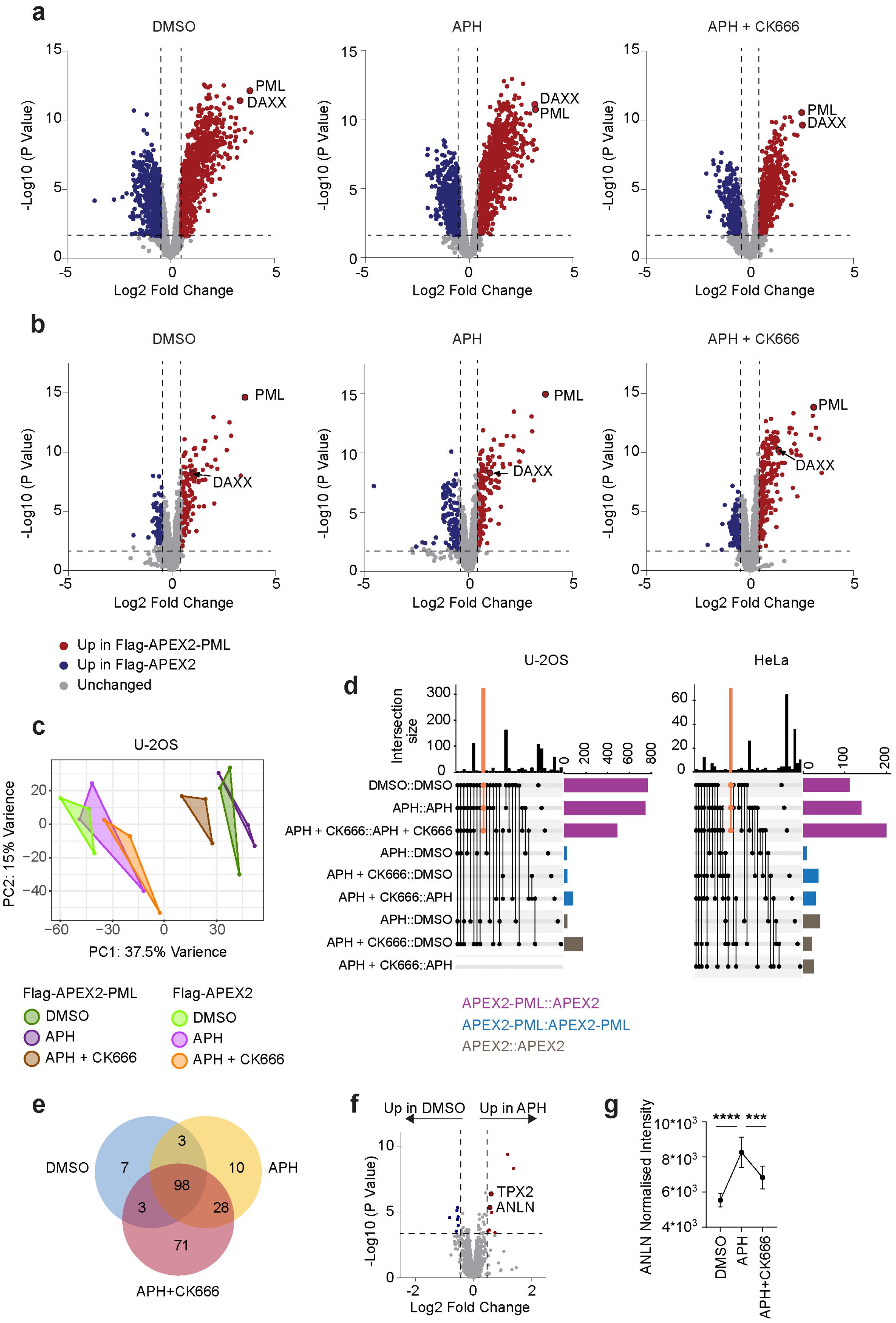

**Supplementary Figure 2 | Validation of the specificity of Flag-APEX2-PML proximity labelling**

**a, b,** Volcano plots showing protein enrichment in Flag-APEX2-PML samples compared to Flag-APEX2 (Empty Vector) in U-2OS (**a**) and HeLa (**b**) cells ($\log_2$ fold change > 0.5, $-\log_{10}$ p-value > 1.3) are highlighted in red; those enriched in Flag-APEX2 controls are shown in blue. Cells were treated for 24 h with DMSO or 0.4 μM aphidicolin (APH), with or without 200 μM CK-666. Statistical analysis: two-sided Student's t-test; n = 3 biological replicates.

**c,** Principal component analysis (PCA) of variance in the biotinylated proteome of U-2OS cells expressing either Flag-APEX2 or Flag-APEX2-PML, treated with DMSO, APH, or APH + CK-666 for 24 h. Each point represents a single biological replicate; n = 3 biological replicates.

**d,** UpSet plots showing the intersection of significantly enriched proteins identified in U-2OS (left) and HeLa (right). Purple bars represent proteins significantly enriched in Flag-APEX2-PML samples relative to Flag-APEX2. Blue and grey denote proteins that vary between treatment conditions within Flag-APEX2-PML and Flag-APEX2 samples, respectively. Horizontal bars indicate the number of proteins shared between the indicated cells and treatments. Orange bars highlight proteins consistently enriched in Flag-APEX2-PML samples across all treatment conditions, representing a core PML NB interactome.

**e,** Venn diagram showing the number and overlap of proteins significantly enriched at PML proximity (enriched at Flag-APEX2-PML vs. Flag-APEX2) in HeLa cells following treatments described in panels (**a, b**).

**f**, Volcano plots showing protein enrichment of Flag-APEX2-PML HeLa cells treated with 24 h of 0.4 μM APH (right) compared to DMSO (left). Significantly enriched PML NB components ($\log_2$ fold change > 0.5, -log p-value > 1.3) are highlighted in red (APH) and blue (DMSO). Statistical analysis: Student's t-test, n = 3 biological replicates.

**g**, Quantification of normalised Anillin (ANLN) intensity from Flag-APEX2-PML HeLa samples under the conditions described in (**a, b**). Statistical analysis by one-way ANOVA followed by Tukey's post hoc test; P < 0.001 (***), P < 0.0001 (****).

#### Supplementary Figure 3

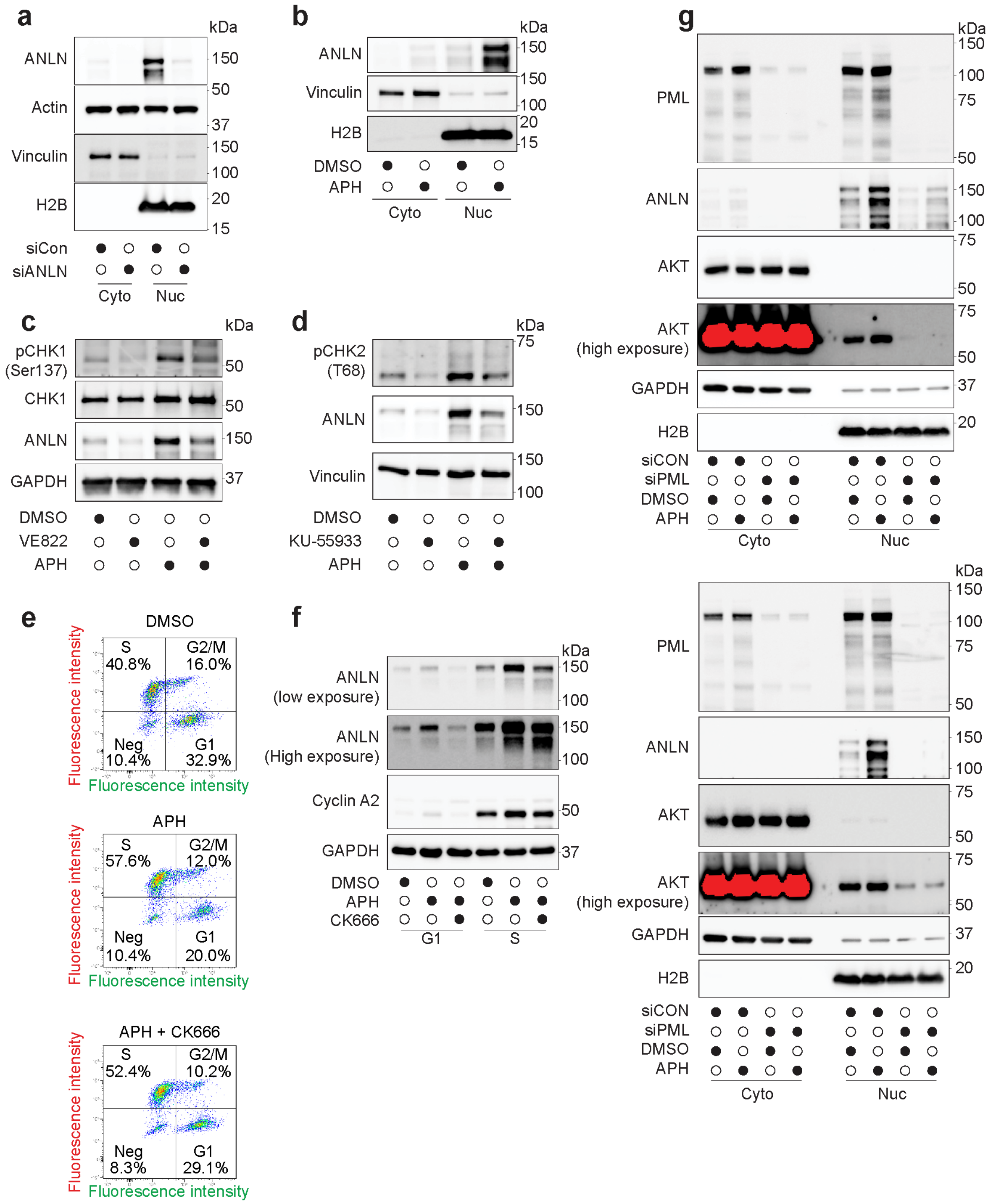

**Supplementary Figure 3 | ATR-, ATM-, and PML-dependent upregulation of nuclear Anillin in response to replication stress**

**a, b,** Representative immunoblots of nuclear and cytoplasmic fractions from U-2OS cells either 72 h post-transfection with the indicated siRNAs (**a**) or treated with 0.4 μM aphidicolin (APH) for 24 h (**b**). Nuclear and cytoplasmic fractions were loaded at a 4:1 volume ratio. n = 3 biological replicates.

**c, d,** Immunoblots of nuclear and cytosolic fractions from U-2OS cells treated with 0.4 μM APH for 24 h in the presence or absence of 0.1 μM ATR inhibitor VE-822 (**c**) or 10 μM ATM inhibitor Ku-55933 (**d**). Nuclear-to-cytoplasmic loading ratio, 4:1. n = 3 biological replicates.

**e,** Scatter plots showing fluorescence-based sorting of Three-colour Fucci U-2OS cells into G1 and S populations following 24 h treatment with DMSO or 0.4 μM APH. n = 3 biological replicates.

**f,** Immunoblots of sorted cell populations from (**e**).

**g,** Two representative immunoblots of nuclear and cytosolic fractions from U-2OS cells transfected with siRNA targeting PML (si-PML) or a non-targeting control (si-Control) for 72 h, followed by treatment with or without 0.4 μM APH for 24 h. Nuclear-to-cytoplasmic loading ratio, 4:1. n = 3 biological replicates.

#### Supplementary Figure 4

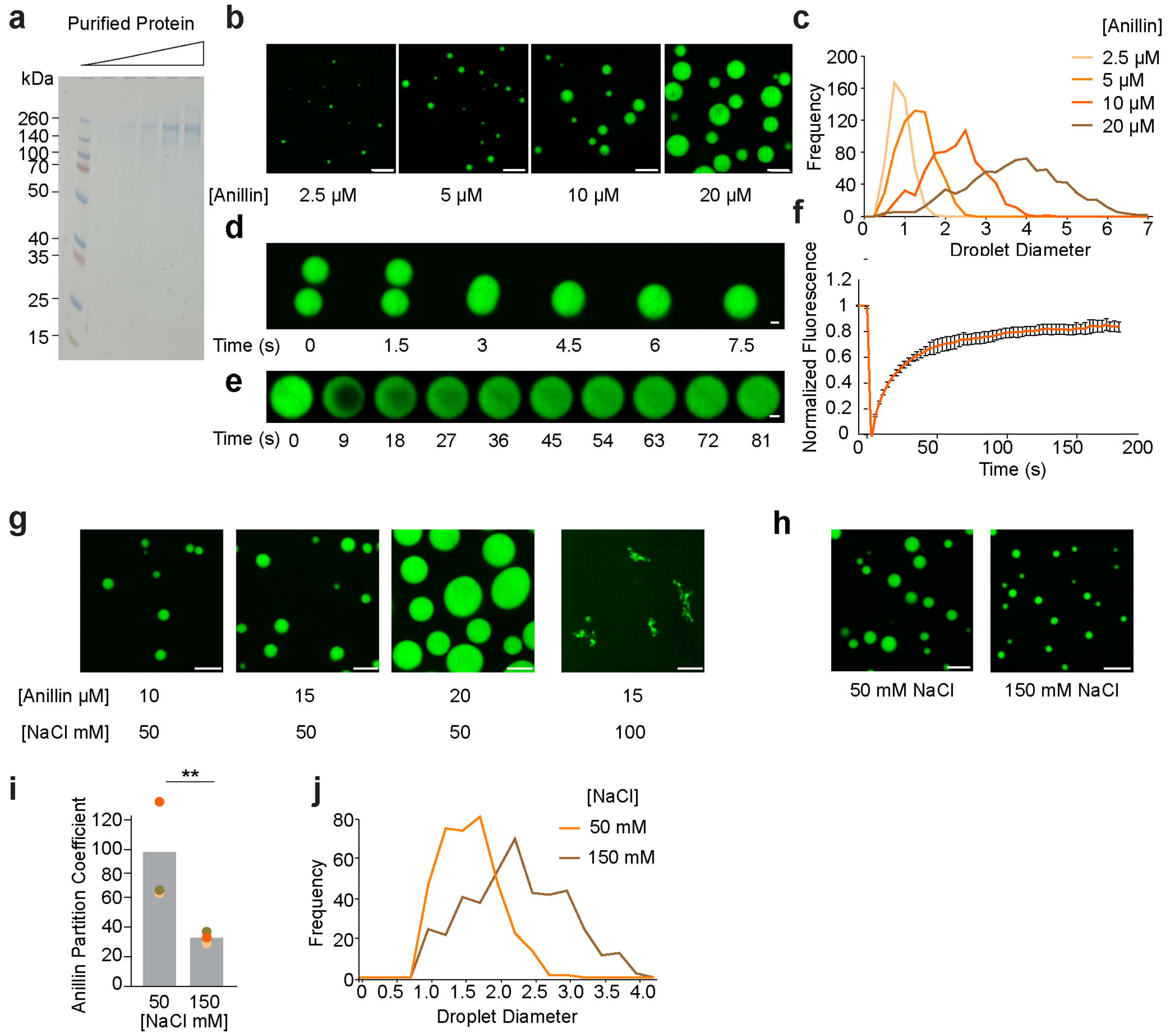

**Supplementary Figure 4 | Anillin phase separates into liquid-like condensates.**

**a,** SDS-PAGE analysis of purified Anillin-GFP with Coomassie staining. n = 3 biological replicates.

**b,** Spinning disk confocal images of Anillin condensates formed *in vitro* at increasing protein concentrations in buffer containing 20 mM Tris (pH 7.4), 150 mM NaCl, 5 mM TCEP, and 3% (w/v) PEG. Scale bar, 5 µm.

**c,** Distribution of condensate diameters under the conditions shown in panel (**b**). n = 3 biological replicates

**d,** Time-lapse spinning disk confocal images showing a fusion event between Anillin condensates (20 µM) in the same buffer conditions. Scale bar, 1 µm.

**e,** Time-lapse spinning disk confocal representative images of fluorescence recovery after photobleaching (FRAP) of an Anillin condensate. Scale bar, 1 µm.

**f,** Quantification of average fluorescence recovery ± s.d. from FRAP analysis of Anillin condensates; n = 19 independent condensates.

**g**, Representative spinning disk confocal images of Anillin condensates formed at increasing protein concentrations in buffer lacking PEG as a molecular crowding agent (Left). Condensates fail to form at 15 µM Anillin in buffers with elevated ionic strength in the absence of PEG (Right). Scale bars, 5 µm.

**h,** Representative confocal images of Anillin condensates formed at 5 µM protein concentration in buffers containing 50 mM or 150 mM NaCl. Scale bars, 5 µm.

**i,** Quantification of Anillin partitioning into condensates under the conditions shown in (**h**). The partition coefficient was calculated as the ratio of fluorescence intensity inside condensates to that in the surrounding solution. Bars represent the mean of three independent experiments; overlaid circles indicate individual replicate means. Statistical analysis by unpaired, two-tailed t-tests; $P < 0.01$ (**).

**e,** Distribution of condensate diameters under the conditions shown in (**h**).

#### Supplementary Figure 5

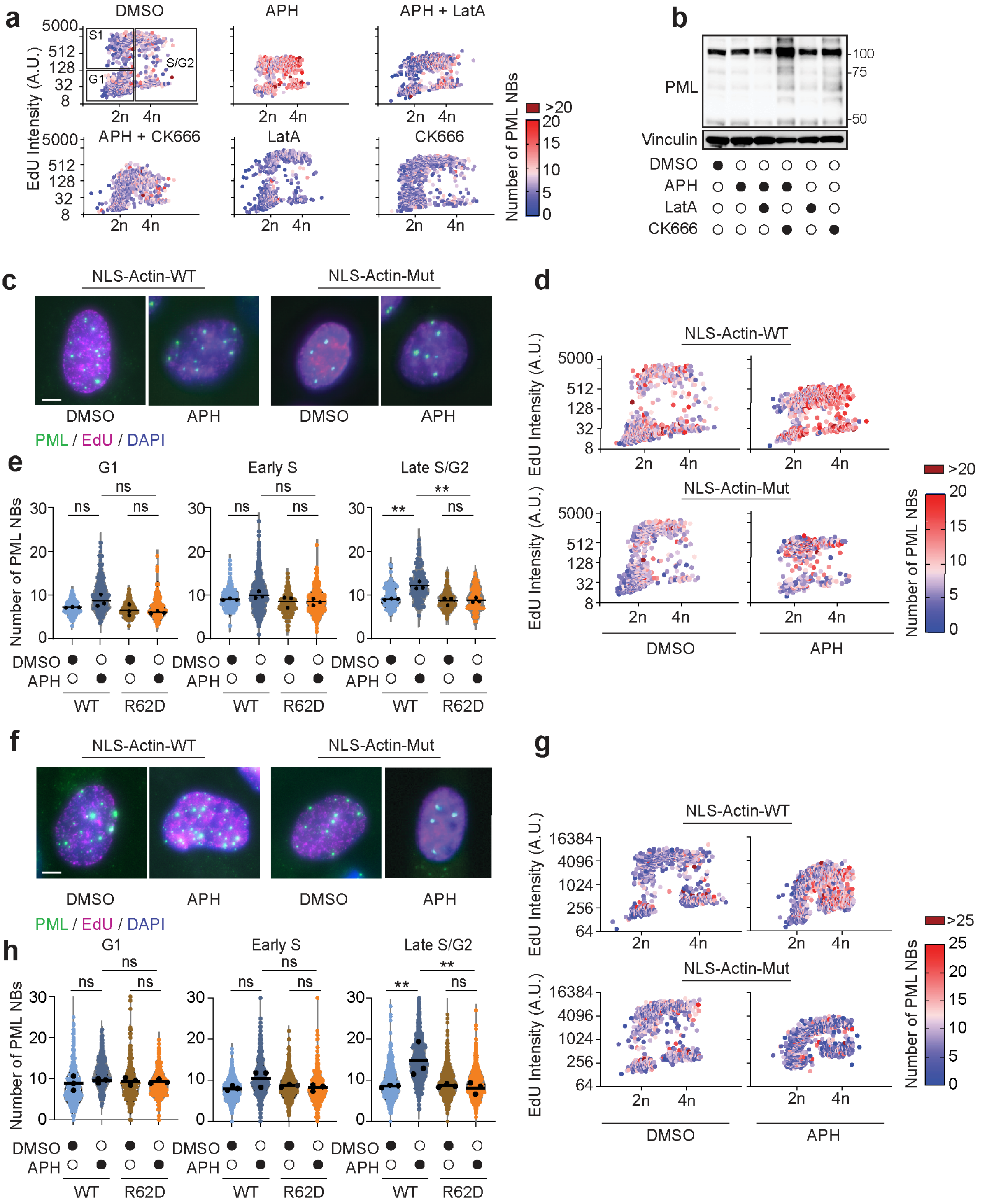

**Supplementary Figure 5 | Nuclear F-actin associates with PML NBs and modulates their condensate behaviour.**

**a,** Scatter plot of individual nuclei showing DAPI versus EdU intensity from HeLa cells pulse-labelled with EdU 1 h before fixation and stained for PML after 24 h treatment with DMSO, 0.4 µM APH, or APH in combination with 200 µM CK-666 or 4 µM latrunculin A (LatA). PML NB count is colour-coded. Black rectangles indicate gates used to define G1, S1 and S2 and G2 populations quantified in (**Fig. 3h**).

**b,** Representative immunoblot of whole-cell lysates from HeLa cells treated as in (**a**). $n = 3$ biological replicates.

**c,** Representative fixed-cell images of U-2OS cells transiently expressing nuclear wild-type actin (NLS-Actin-WT) or a polymerisation-deficient mutant (NLS-Actin-Mut). Cells were pulse-labelled with EdU for 1 h before fixation and stained for PML after 24 h DMSO or 0.4 µM APH. Scale bar, 5 µm.

**d,** Scatter plots of nuclei from (**c**), showing DAPI intensity versus EdU incorporation, colour-coded by PML NB number.

**e,** Violin plots of PML NB number across cell-cycle phases from (**c**, **d**). Black dots indicate the average of three independent repeats; solid lines show the mean across replicates. Statistical analysis by one-way ANOVA followed by Tukey's post hoc test; Not significant (ns), $P < 0.01$ (**).

**f,** Representative fixed-cell images of HeLa cells treated as in (**c**). Scale bar, 5 µm.

**g,** Scatter plots of nuclei from (**f**), as in (**d**).

**h,** Violin plots of PML NB number from (**f**, **g**), as in (**e**). Statistical analysis as in (**e**).

#### Supplementary Figure 6

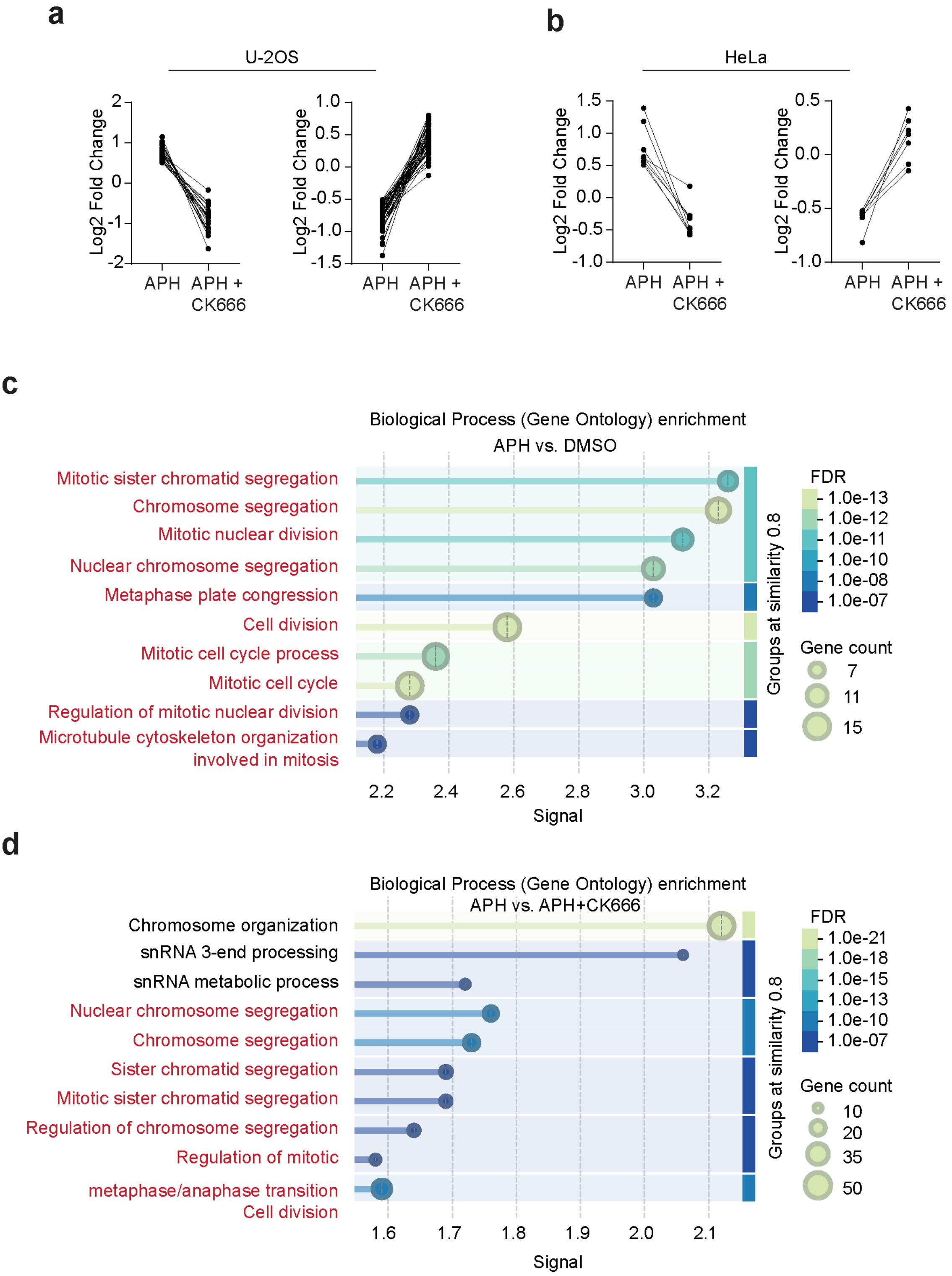

**Supplementary Figure 6 | Nuclear F-actin modulates PML NB's interactome.**

**a, b,** Quantification of biotinylated proteins enriched in Flag-APEX2-PML U-2OS (**a**) and HeLa (**b**) cells treated for 24 h with DMSO, 0.4 μM APH, or APH combined with 200 μM CK-666. Each dot represents one significantly altered protein. In the "APH" column, protein enrichment (left) or depletion (right) is shown relative to DMSO. In the "APH+CK-666" column, the same proteins were plotted relative to APH alone; connecting lines indicate that CK-666 reverses the APH-induced effect.

**c, d,** STRING clustering of proteins from (**a**) enriched at PML NBs after APH versus DMSO (**c**) or depleted in APH+CK-666 versus APH (**d**). Pathways in red font correspond to mitotic processes.

164 **Supplementary Figure 7**

165

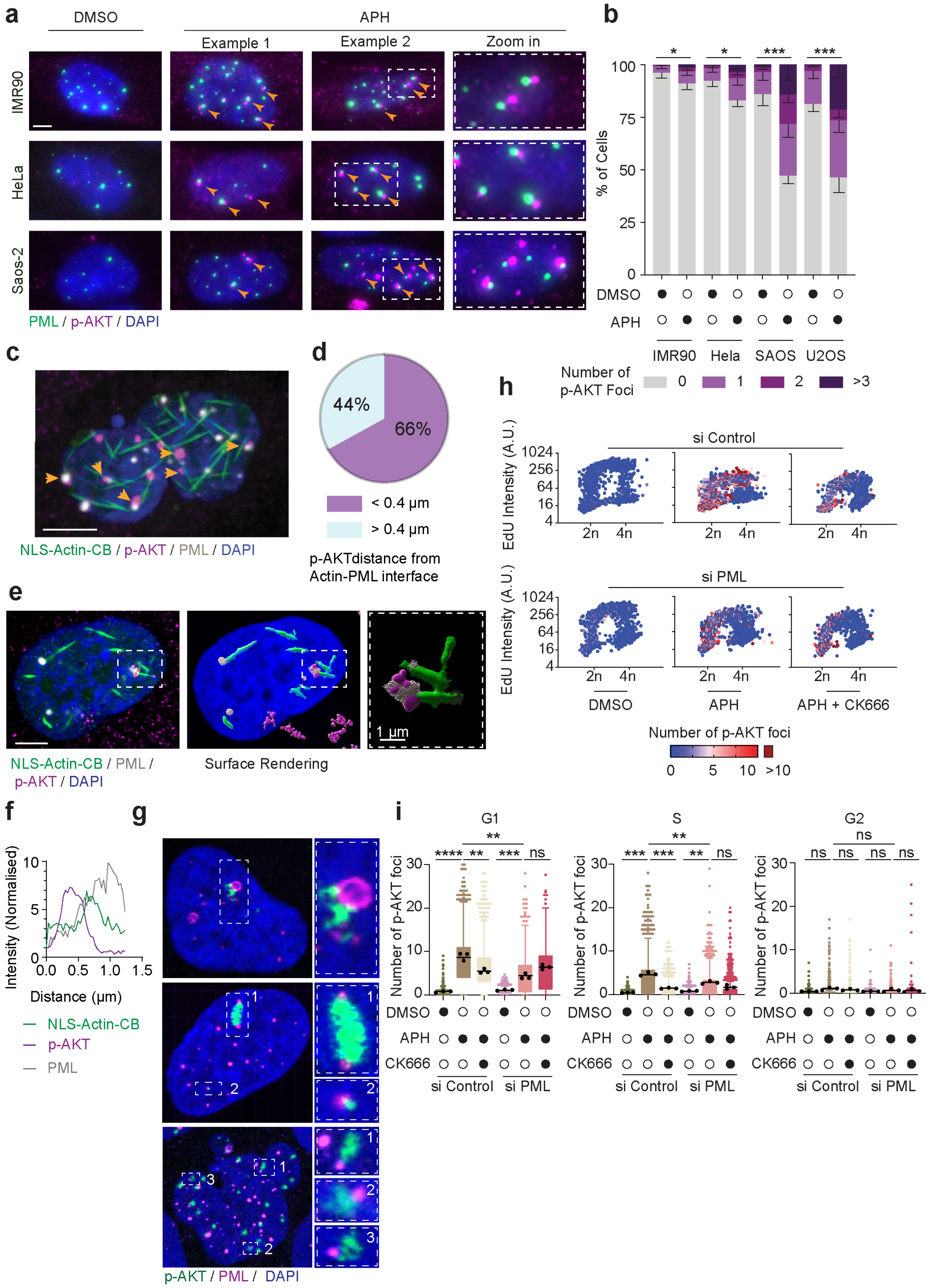

**Supplementary Figure 7 | Nuclear F-actin-PML NB association is required for AKT phosphorylation.**

**a,** Immunofluorescence of IMR90, HeLa, and Saos-2 cells stained for PML and p-AKT-Ser473 after 24 h of 0.4 µM APH treatment. Insets and orange arrowheads highlight p-AKT puncta adjacent to PML NBs. Scale bar, 5 µm.

**b,** Quantification of p-AKT-Ser473 foci per nucleus in the indicated cell lines from (**a**) after DMSO or APH treatment. Statistical analysis by Fisher's exact test; *P < 0.05; **P < 0.01.

**c, e,** Super-resolution Airyscan images (single Z-planes) of U-2OS cells expressing NLS-actin CB and treated with APH for 24 h, stained for p-AKT-Ser472 and PML. Insets and arrowheads highlight p-AKT signal appearing to be released from PML NBs at the tip of nuclear actin filaments. Scale bar, 5 µm.

**d,** Quantification of p-AKT release events associated with nuclear F-actin from (**c**). A total of 342 events were scored across 52 nuclei from 3 independent experiments.

**f,** Fluorescence intensity profile along the dashed line in (**e**), showing nuclear F-actin signal positioned between p-AKT and PML, consistent with an actin-guided release interface.

**g,** Airyscan image of U-2OS cells treated with APH for 24 h and stained for p-AKT-Ser473 and PML. Insets show diffuse p-AKT surrounding PML NBs, consistent with localised release. Scale bar, 5 µm.

**h,** Scatter plot of nuclei from U-2OS cells transfected with si-Control or si-PML for 72 h, pulse-labelled with EdU for 1 h, and stained for p-AKT-Ser473 after DMSO, APH, or APH+CK-666 treatment. Each point represents a nucleus, colour-coded by p-AKT foci number.

**i,** Violin plots of p-AKT foci across cell-cycle phases from (**h**). Black dots represent averages of three independent replicates; solid lines show overall means. Statistical analysis by one-way ANOVA followed by Tukey's post hoc test; Not significant (ns), P < 0.01 (**), P < 0.001 (***), P < 0.0001 (****).

192 **Supplementary Figure 8**

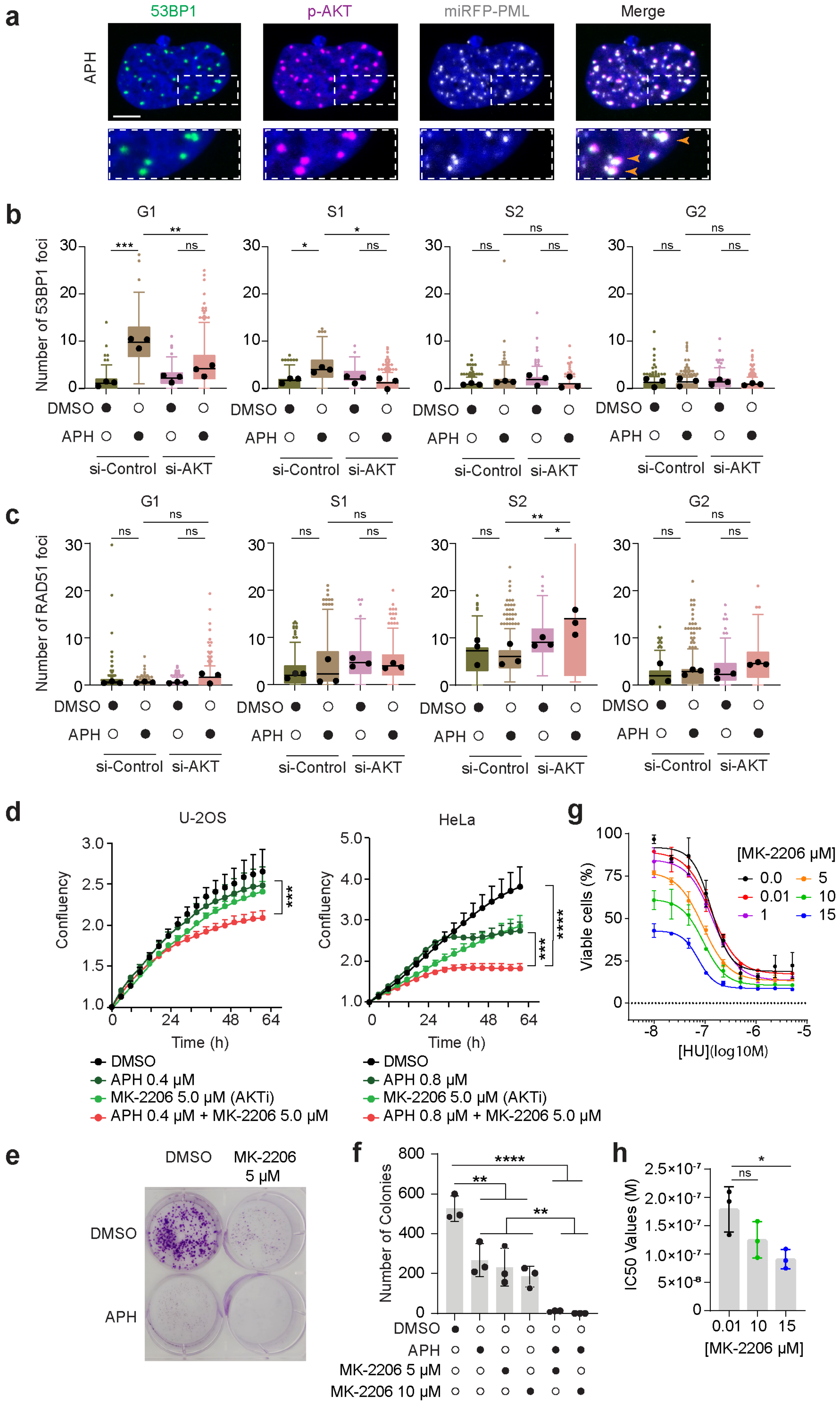

**Supplementary Figure 8 | AKT promotes survival in response to replication stress.**

**a,** Airyscan microscopy (single Z-plane) of miRFP-PML-U-2OS cells treated with 0.4 µM APH for 24 h, stained for p-AKT-Ser473 and 53BP1. Scale bar, 5 µm.

**b, c,** Box plots of 53BP1 (**b**) and RAD51 (**c**) foci number across cell-cycle phases in U-2OS cells transfected with si-Control or si-AKT for 72 h, pulse-labelled with EdU for 1 h, and stained after 24 h of DMSO or 0.4 µM APH treatment. Box plots show values from three biological repeats; black dots indicate replicate means and black lines overall means. Statistical analysis by one-way ANOVA followed by Tukey's post hoc test; $P < 0.05$ (*), $P < 0.01$ (**), $P < 0.001$ (***).

**d,** Normalised confluency curves of U-2OS (left) and HeLa (right) cells treated with 0.4 or 0.8 µM APH, 5 µM AKT inhibitor MK-2206, or combinations thereof (mean ± SD; n = 3). Statistical analysis by one-way ANOVA followed by Tukey's post hoc test; $P < 0.01$ (**), $P < 0.001$ (***), $P < 0.0001$ (****).

**e,** Representative colony formation assay of U-2OS cells treated with 0.4 µM APH, 5 µM or 10 µM MK-2206, or combinations.

**f,** Quantification of colonies from (**e**). Bars show the mean of three independent repeats; black dots indicate replicate means. Statistical analysis by one-way ANOVA followed by Tukey's post hoc test; $P < 0.05$ (*), $P < 0.01$ (**), $P < 0.001$ (***), $P < 0.0001$ (****).

**g,** Hydroxyurea (HU) dose-response curves of U-2OS cells treated with the indicated concentrations of MK-2206 for 24 h.

**h,** $IC_{50}$ values from (**g**). Bars show mean of three biological repeats; black dots indicate individual replicates. Statistical analysis by unpaired, two-tailed t-tests; Not significant (ns), $P < 0.05$ (*).

#### Supplementary Figure 9

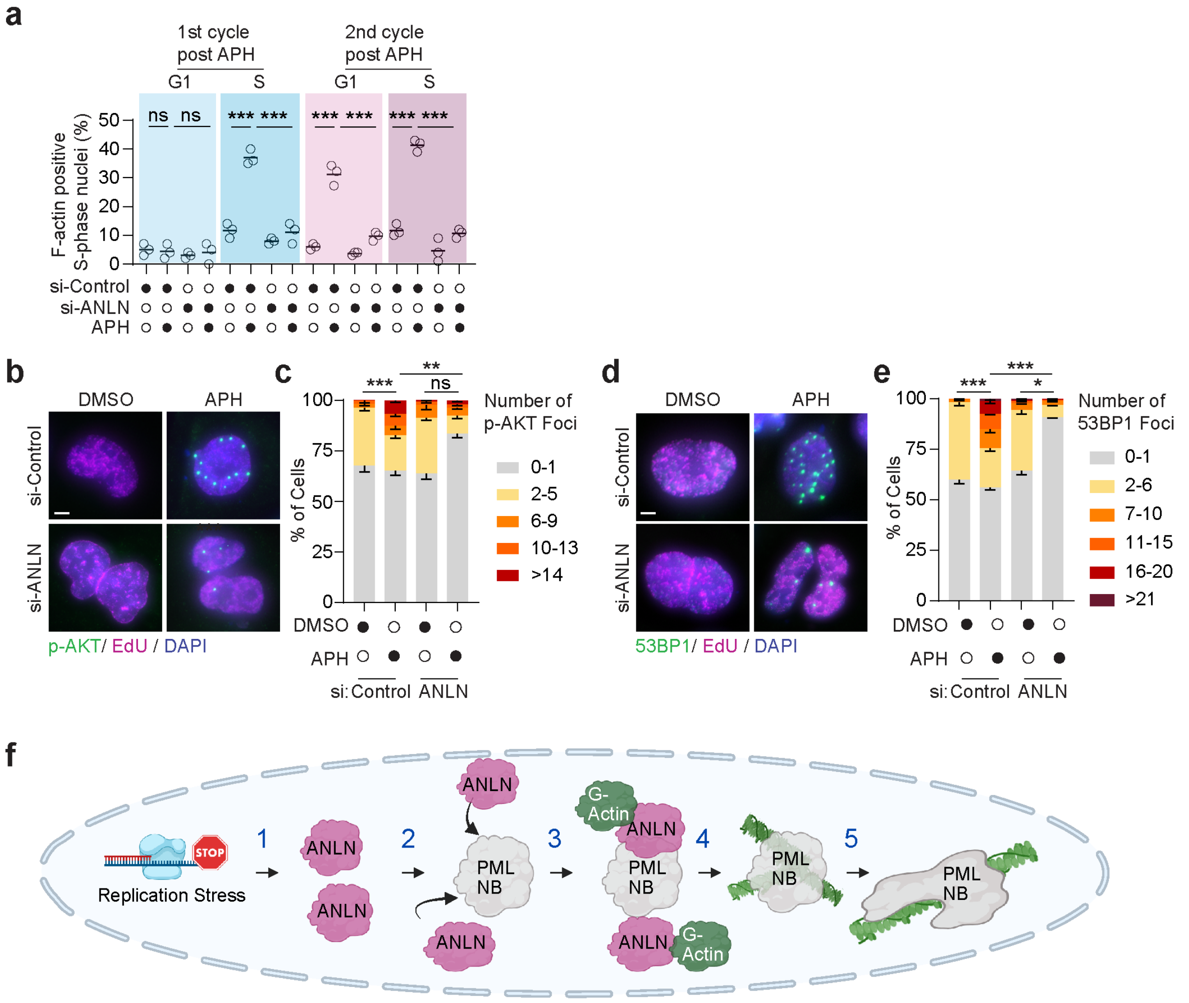

**Supplementary Figure 9 | Anillin is required to protect under-replicated DNA.**

**a,** Quantification of G1 and S-phase nuclei positive for nuclear F-actin. Cells were transfected with the indicated siRNAs 48 h before imaging and treated with DMSO or APH at the start of imaging for a total of 96 h. Data are shown as mean ± s.e.m.; n = 3 biological replicates, ≥32 nuclei per replicate. Statistical analysis by two-sided Fisher's exact test; Not significant (ns), $P < 0.05$ (*), $P < 0.01$ (**), $P < 0.001$ (***).

**b, d,** Representative fixed-cell images of U-2OS cells transfected with si-Control or si-Anillin 72 h before fixation, pulse-labelled with EdU for 1 h, and stained for p-AKT-Ser473 (**b**) or 53BP1 (**d**) after 24 h of DMSO or 0.4 µM APH treatment. Scale bar, 5 µm.

**c, e,** Quantification of p-AKT (**c**) or 53BP1 (**e**) foci from the experiments in (**b, d,** respectively). Bar graphs show mean ± s.e.m.; n = 3 biological replicates. Statistical analysis by one-way ANOVA followed by Tukey's post hoc test; $P < 0.05$ (*), $P < 0.01$ (**), $P < 0.001$ (***), $P < 0.0001$ (****).

**f,** Model for Anillin-dependent nuclear actin assembly at PML NBs in the replication stress response. (1) Replication stress induces increased nuclear Anillin (ANLN); (2) Anillin accumulates at PML NBs; (3) Local enrichment of Anillin drives G-actin partitioning at PML NBs; (4) Concentrated G-actin spontaneously polymerises into filaments; (5) Growing filaments generate mechanical forces that deform PML NB condensates.

**Supplementary Movie 1 | Nuclear actin assembles at PML NBs in response to replication stress.**

Representative maximum projection movie acquired by spinning-disk confocal live-cell microscopy of a U-2OS cell co-expressing nuclear-actin-CB, PCNA-CB, and stably expressing miRFP-647-PML. Cells were transfected 48 h prior to imaging and treated with 0.4 μM APH at the start of the imaging session. Time is shown as hours:minutes:seconds relative to the first frame. Nuclear registration (to stabilise the nucleus at the frame of origin) was performed as previously described[3]. A slow-motion annotated segment highlights the early stages of nuclear actin assembly at PML NBs.

**Supplementary Movie 2 | Nuclear actin assembles at PML NBs in response to replication stress.**

Maximum projection spinning-disk confocal live-cell movie of a HeLa cell expressing nuclear-actin-CB and stably expressing miRFP-647-PML, treated with 0.4 μM APH. Cells were transfected 48 h before imaging, and APH was added at the start of the imaging session. Time is displayed as minutes:seconds relative to the first frame.

**Supplementary Movie 3 | Wetting-like behaviour of PML NBs and nuclear F-actin.**

Representative maximum projection spinning-disk confocal live-cell movie of a U-2OS cell expressing nuclear-actin-CB, PCNA-CB, and stably expressing miRFP-647-PML. Cells were treated as in Supplementary Movie 1 (0.4 μM APH; transfected 48 h before imaging). Time is displayed as hours:minutes:seconds relative to the first frame. Nuclear registration was performed as previously described[3].

**Supplementary Movie 4 | Wetting-like behaviour of PML NBs and nuclear F-actin.**

Representative maximum projection spinning-disk confocal live-cell movie of a HeLa cell expressing nuclear-actin-CB and stably expressing miRFP-647-PML. Cells were treated as in Supplementary Movie 2 (0.4 μM APH; transfected 48 h before imaging). Time is displayed as hours:minutes:seconds relative to the first frame.

**Supplementary Movie 5 | F-actin maintains PML NBs in a dispersed state.**

Representative maximum-projection spinning-disk confocal live-cell movie of a HeLa cell expressing nuclear-actin-CB and stably expressing miRFP-647-PML, treated with 0.4 μM APH as in Supplementary Movie 2. Time is shown as hours:minutes:seconds relative to the first frame. Nuclear registration was performed as previously described[3].

**Supplementary Movie 6 | Nuclear AKT activity positively correlates with actin assembly.**

Representative maximum-projection spinning-disk confocal live-cell movie of a U-2OS cell expressing nuclear-actin-CB and stably expressing FoxO1-mStrawberry, treated with 0.4 µM APH at the start of the imaging session. Dual-channel and individual-channel views are shown side by side. A slow-motion annotated segment highlights the decline in AKT activity coinciding with the onset of nuclear actin assembly. Time is displayed as hours:minutes:seconds relative to the first frame.
